# Supracellular Mechanics and Counter-Rotational Bilateral Flows Orchestrate Posterior Morphogenesis

**DOI:** 10.1101/2025.11.18.689090

**Authors:** Geneva Masak, Lance A Davidson

## Abstract

After gastrulation, the tailbud of Xenopus laevis emerges as a morphogenetic engine. We find that the transition from late neural to early tailbud stages is characterized by large-scale, counter-rotational tissue flows flanking the blastopore, and spatially coupled to dorsal elongation and ventral compression. Live imaging and quantitative flow analysis reveal that these rotations are maintained over ∼200 µm, suggesting long-range mechanical coupling between dorsal and ventral tissues. High-resolution confocal microscopy shows fibronectin and laminin fibrils radiating ventrally from the blastopore in spoke-like arrays between ectoderm and mesoderm. Perturbations of cell proliferation, radial intercalation, and mediolateral intercalation fail to abolish these flows, whereas targeted disruption of ventral extracellular matrix integrity severely impairs rotational movement. These results identify a previously unrecognized ventral ECM network as a critical mechanical scaffold in regulating posterior tissue rotation, highlighting the interplay between ECM organization, tissue mechanics, and morphogenetic flow during tailbud development.

## Introduction

Large-scale tissue flows of gastrulation and neurulation reshape the early embryo from a spherical mass into a tube-like structure, laying the foundation for the formation of the central nervous system and organs. These movements requires tissues to act as cohesive, mechanically integrated structures. For instance, primitive streak formation in avians involves large scale cohesive movements as the epiblast moves toward the midline and simultaneously undergoes lateral displacement, generating broad, organized rotations within the epithelial sheet (Henkels et al., 2013).This combination of converging and shearing movements produces rotational flow patterns or “vortices”, where adjacent tissues dynamically adjust to accommodate streak elongation (Henkels et al., 2013). Such rotational flow patterns have been seen in during formation of hair follicles in mouse (Cetera et al., 2018) and presomitic mesoderm in fish (Banavar et al., 2021; Stooke-Vaughan et al., 2025). Similarly, neural tube formation relies on a series of organized movements that transform the flat neural plate into a cylindrical structure (Davidson and Keller, 1999; Lowery and Sive, 2004; Kinoshita et al., 2008). This process begins with the elevation and mediolateral convergence of the neural folds and culminates with tube closure (Morita et al., 2012). During neurulation, dorsal ectoderm, mesoderm, and endoderm all extend along the anterior posterior axis. These two examples illustrate how mechanical processes in multiple tissues are integrated to shape the embryo.

Morphogenetic movements are often attributed to forces generated by discrete cell behaviors such as cell division, radial intercalation, and directed cell rearrangement (Davidson, 2024). Cell division can be a source for tissue growth (His, 1888) with oriented cell division contributing to posterior axis elongation (Firmino et al., 2016). Radial intercalation can alter tissue topology and structural organization by redistributing cellular volume from one layer to another (Davidson and Keller, 1999; Stubbs et al., 2006; Rauzi, 2020; Ventura and Sedzinski, 2022). Similarly, directed cell rearrangement, i.e., cell intercalation within a tissue plane, can drive large scale convergent extension movements (Keller, 2002). Long-range physical constraints provided by a polarized myosin II distribution can also convert rearrangement behaviors into rotational flows (Sato et al., 2015). Thus, a diverse array of cell behaviors can contribute to precise morphological changes required during development.

The cytoskeleton and ECM are thought to coordinate large-scale tissue movements (Loganathan et al., 2016), however, their contributions to rotational tissue movements remain poorly understood. Actin filaments, microtubules, and intermediate filaments provide structural support and facilitate coordinated cell movements, with actomyosin contractility responsible for the forces that drive convergent extension and tissue rotation in both vertebrate and invertebrate development (Zamir et al., 2008). Similarly, integrin cell surface receptors anchor cells to the ECM and transmit forces into cells and across tissues. The ECM provides both structural support and mechanical cues that regulate cell migration and tissue flow, with fibronectin and collagen fibers aligning with the direction of tissue elongation (Rozario and DeSimone, 2010). ECM stiffness gradients may also serve as physical constraints that modulate the movement and positioning of cells, a factor that could be relevant in the establishment of counter-rotational flows (Gjorevski and Nelson, 2010).

In this paper we describe counter-rotational flows lateral to the blastopore, ventral tissue compaction, and dorsal convergent extension in the aquatic frog *Xenopus laevis* during the transition between neural tube closure and tailbud morphogenesis. We quantify deformation and rotational flows from time-lapse image sequences and test the contributions of cell division, radial cell intercalation, ECM assembly, tissue patterning, and dorsal convergent extension in generating counter-rotating vortices. Our findings suggest that posterior morphogenesis and counter rotational lateral flows are the result of long-range, externally driven forces constrained by ventral ECM.

## Results

### Emergence of Counter-Rotational Tissue Flows in the Posterior Embryo

We first examined the development of the blastopore and surrounding tissues from late neural to early tailbud (stages 18 to 24, NF; see lateral view, Fig 1A)(Beck, 2015). Changes in the posterior domain from stages 18 to 20 are subtle in contrast to cranial closure and progressive pigmentation of the cement gland (see anterior view, Fig 1A). Lateral views from stage 20 to 24 reveal narrowing and lengthening along the anteroposterior axis and ventral movement of the blastopore (lateral view, Fig 1A).

**Figure 1.**
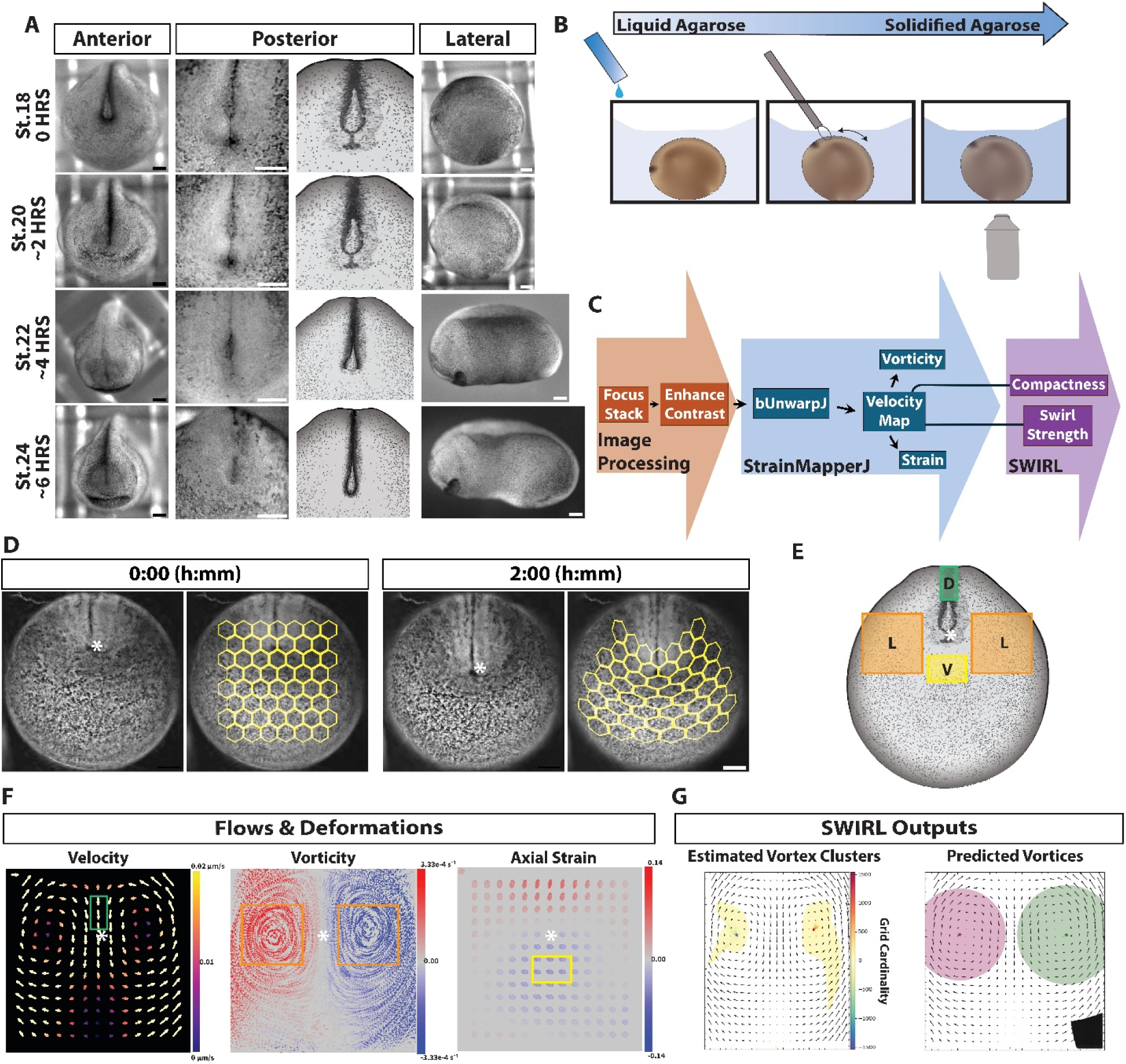
Static and kinematic analysis of the blastopore and posterior tissues. (A) Representative images of the anterior, posterior, and lateral view of Xenopus embryos from late neural to early tailbud stages with schematic of posterior view. (B) Embedding embryos in agarose for posterior imaging. (C) Image analysis pipeline. (D-F; blastopore, *). (D) Brightfield images from stage 18 to stage 20. Hexagonal mesh deformation maps overlaid in yellow, illustrating cumulative tissue displacement (see also Video S1). (E) Regions of interest around the blastopore (green, dorsal; orange, lateral; yellow, ventral). (F) Maps of tissue flow and deformation. Left: velocity vector map (direction, see Fig S1A; magnitude, see Fig S1B). Middle: vorticity map (Fig S1B) displaying rotational motion, superimposed onto randomized dot plots deformed by displacement and max-projected across all timepoints to represent movement patterns (Fig S1A). Right: Axial strain map represented as ellipse maps (Fig S1A) indicating principal strain direction and magnitude (Fig S1B). (G) Vortex detection using SWIRL. Estimated vortex centers are represented by clusters of grid cardinality values overlaid onto velocity vector maps. Pink and green circles indicate the predicted vortex size and location, while stars mark the corresponding vortex centers. (A, D) scale bars, 200 µm.

To better understand these dynamics, we turned to quantitative analysis of brightfield timelapse image sequences (Fig 1B). To quantify posterior tissue deformations, we utilized a custom image analysis pipeline to generate projected 2D maps of deformation, velocity, elastic strain, and vorticity (Fig 1C). To aid in visualizing deformation, we overlaid a yellow hexagon mesh onto the tissue surface; distortions of the mesh pattern indicate the magnitude and direction of tissue deformation and a velocity map (Fig 1D; See Video S1). We use the blastopore as a reference point for assessing adjacent tissue movements, and describe tissues as dorsal, ventral, or lateral to reference their position relative to the blastopore (Fig 1E). These three regions reveal distinct spatial patterns of deformation (Fig 1E). The dorsal region, comprised of neural tissues directly dorsal to the blastopore, displayed elongation and narrowing of axial tissues, consistent with known convergent extension movements (Fig 1D-F)(Keller and Sutherland, 2020). The ventral ectoderm, located approximately 100 µm ventral to blastopore exhibited movements opposite of dorsal tissues, with midline cells at the midpoint flowing laterally, while undergoing axial compression (Fig 1D-F). Meanwhile, the lateral regions, which encompasses the ectoderm positioned on either side of the blastopore undergo counter-rotational movements similar to “polonaise flow” observed during avian gastrulation (Wang and White, 2021) and adjacent to the presomitic mesoderm in the zebrafish tailbud (Stooke-Vaughan et al., 2025). These Velocity maps highlight distinct directional motion in dorsal, lateral, and ventral regions; velocities in dorsal regions exceeded 0.02 µm/s while ventral velocities were near zero (Fig 1F; Fig S1A-B). In the lateral regions, velocities also approached zero near the midline but increased, reaching peak values at the periphery of the counter-rotational domains (Fig 1F; Fig S1A-B).

From these initial assessments we focused on motions in lateral and ventral regions. Velocity fields describe the magnitude and direction of tissue displacement, but they do not isolate the rotational component of motion. Given the presence of counter-rotational movements lateral to the blastopore, we calculated vorticity, a measure that indicates local rotation of the flow (see Supplemental Material S1; (Saadaoui et al., 2020; Asai et al., 2024)). Vorticity maps reveal two counter-rotating flow regions that are sharply delineated at the midline where they are separated by the blastopore (Fig 1F; Fig S1A-B). Since we observed hexagonal cells in the ventral region of the deformation mesh shortening and widening (Fig 1D), we calculated axial strain (see Document S1). Axial strain maps revealed a sharp transition between dorsal and ventral tissues, with strain values reaching zero across the blastopore (Fig 1F; Fig S1B). Dorsal tissues exhibited positive strain (0.14 per hour) consistent with axial elongation, while ventral tissues showed negative strain (-0.14 per hour), indicating compaction of the region (Fig 1F). This strain transition around the blastopore suggests differential mechanical loading in these regions where movement is redirected rather than halted (Fig 1F). In addition to dorsal-ventral differences, lateral regions displayed a gradient of axial strain across the counter-rotations, suggesting these motions were driven by shear forces (Fig 1F). Since vorticity does not differentiate between stable vortices and shear-driven deformation (Tian et al., 2018) we sought a more robust method to distinguish true rotational structures from shear-driven artifacts. Using SWIRL analysis (see Supplemental Material S1), we identified multiple stable and spatially restricted vortices within the posterior tissue (Fig 1G). Swirling strength and compactness (Fig S1C) suggest that these counter-rotating structures are both persistent and spatially constrained, rather than transient shear-driven flows.

In biophysical, geophysical, and astrophysical systems, rotational flows can arise through two primary mechanisms: free vortex motion, in which rotation results from locally generated forces, or forced vortex motion, where rotation is driven by externally generated forces together with global constraints on movement (Sinha et al., 2013). A defining feature of these two vortex types is the relationship between velocity and radial distance from the vortex center. In a free vortex, tangential velocity decreases with distance, while in a forced vortex, velocity increases with distance due to coordinated motion across the system. Boundaries, such as tissue interfaces or ECM networks, often determine which type emerges and how flows are maintained (Davidson, 2011). In embryonic tissues, however, these flows ultimately depend on the collective behaviors of individual cells: their polarity, rearrangements, and interactions with the extracellular environment (Wang et al., 2020; Wu et al., 2023). Thus, distinguishing between free and forced vortex dynamics provides a framework for interpreting posterior flows, but also raises a key question about how the behaviors of individual cells might contribute to the emergence and maintenance of these large-scale vortices.

### Nuclear Trajectories and Cell Rearrangement Reveal Dynamic Tissue Mechanics

To directly assess how individual cell behaviors contribute to large-scale vortex formation, we used live confocal timelapse imaging to track nuclei. Nuclei trajectories remained consistent with global patterns identified in brightfield images. Nuclei in the dorsal regions exhibiting the highest velocities, while cells in the ventral region display velocities close to zero (Fig 2A; See Video S2). We analyzed the relationship between average nuclear track velocity and radial distance from the vortex center and found a positive correlation (<r²> = 0.33) (Fig 2B), consistent with forced vortex flow (Sinha et al., 2013). This suggests that counter rotations in the embryo are driven by externally generated forces, rather than emerging from active cell migration or intrinsic force generation within the rotating regions. Additionally, the low velocities of cells in ventral regions do not appear to impede cells in dorsal and lateral regions, instead resembling a dynamic boundary, where movements diverge bilaterally rather than continuing posteriorly (Fig 2A).

**Figure 2.**
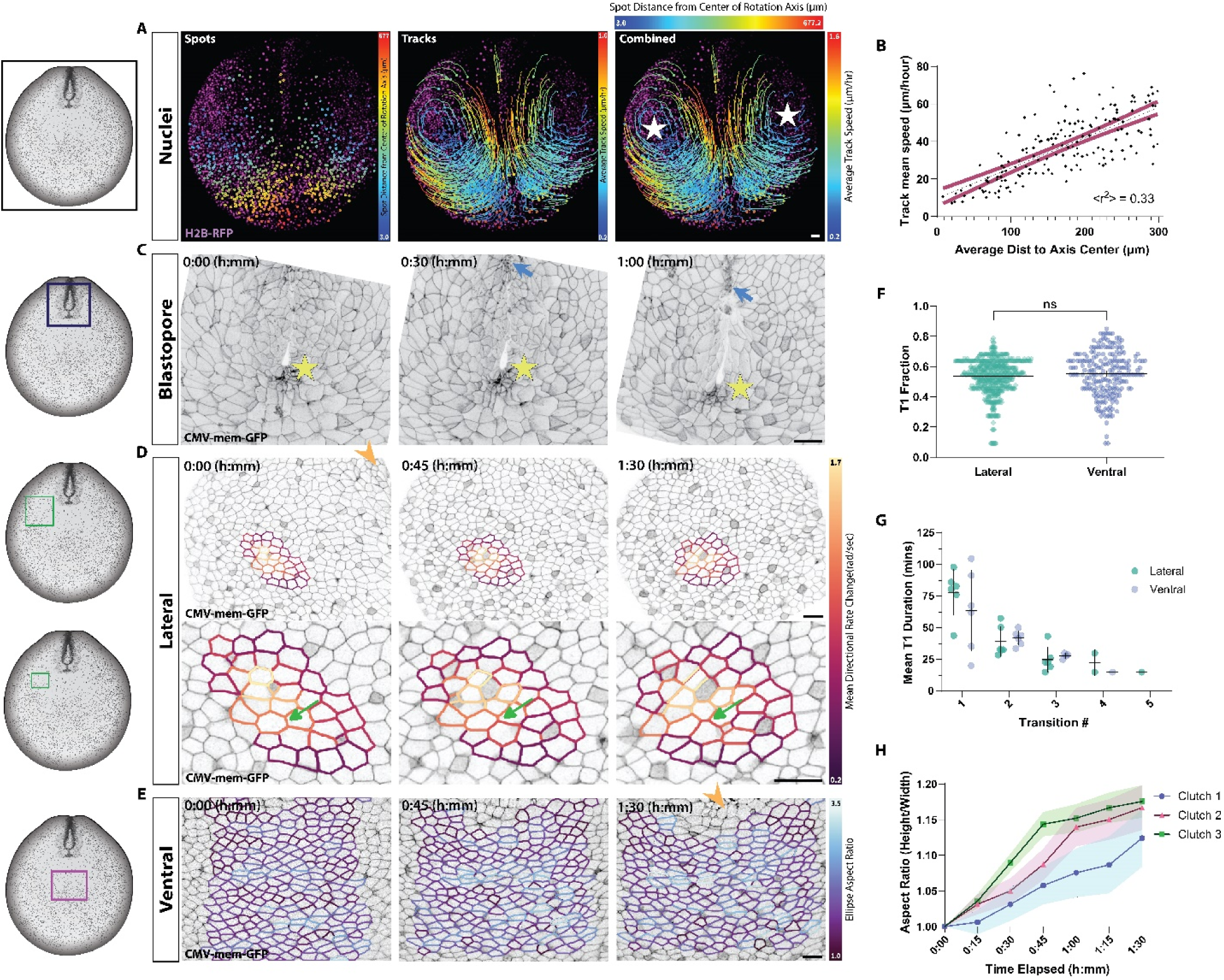
Live confocal imaging of nuclear and membrane-labeled embryos reveals distinct cell and tissue dynamics. (A,C-E) Schematics of stage 18 embryo indicating approximate region of imaging (box) relative to the blastopore, captured in 6 embryos from 3 separate clutches. (A) Nuclear trajectories over 3 hours in an H2B-labeled embryo (see also Video S2). Dot color indicates distance to center of rotation (white stars); track colors indicate average velocity. (B) Nuclear velocity plotted as a function of radial distance from the vortex center for the representative embryo shown in (A) (169 nuclei). R² reflects the mean goodness-of-fit across 6 embryos (Simple linear regression). Black dotted line, linear regression fit; red shading, 95% CI. (C-E) Frames for representative timelapse of mem-GFP transgenic embryos. (C) Tissue shape changes at 30 min. intervals (see also Video S3). Cells below the opening of the blastopore constrict (yellow star) as zippering neural cells converge (blue arrow). (D) Clockwise-rotating lateral tissues. A representative subset of tracked cells is outlined and colored based on their mean directional rate change (radians/sec). Note: the blastopore (orange arrowhead) moves out of view as rotation progresses. T1 transitions (green rounded arrow) occur throughout tissue (see also Video S4). (E) Cell shape changes within ventral tissues. Tracked cells are outlined and color-coded by aspect ratio. Note: the blastopore (orange arrowhead) enters the field of view in the final frame (see also Video S5). (F) Fraction of time spent in a T1 transition by individual cells in lateral and ventral tissues. Symbols represent individual cells pooled across 6 embryos per region; per-embryo cell counts ranged from 27 to 528 (Mann-Whitney U, p=0.72). (G) Duration of T1 transitions. Time cells spent in each T1 transition in lateral and ventral tissues. Symbols represent the mean per embryo. (H) Normalized cell aspect ratios. Colors represent the average within a clutch. Symbols represent mean per embryo, with 78–450 cells analyzed in each embryo. Colored area fill; 95% CI. (A,C-E) scale bars 50 µm.

We next examined how the blastopore remodels, given its distinct position between these counter-rotational flows. Timelapse confocal imaging of *CMV-mem-GFP* embryos reveals a more detailed view of the circumblastoporal collar with blastopore narrowing as posterior neural folds fuse (Fig 2C; See Video S3). As previously reported, we observed a transient release of extracellular fluid (Kato and Inomata, 2023), temporarily expanding the blastopore before it narrowed again (data not shown). Additionally, cells exhibited visible constriction just within the blastopore lip, coinciding with blastopore narrowing (Fig 2C; yellow star). These observations suggest a complex set of forces and deformations occur around the blastopore, possibly coupling dorsal, ventral or lateral tissues. Since large-scale tissue flows are commonly ascribed to collective behaviors of individual cells, we next examined cell morphology and movement patterns within ventral and lateral regions. Time-lapse confocal imaging of membrane-labeled transgenic embryos revealed dynamic cell behaviors, including cell division, radial intercalation, and convergent extension (Fig 2D-E; See Video S4-5). These processes were present in both ventral and lateral regions; however, the strong rotations observed in lateral regions suggested the possibility of more frequent cell rearrangements in these areas.

To quantify cell rearrangement dynamics we tracked T1 transitions (Fig S2A-C; see Supplemental Material S1). The median T1 dwell fraction was 41% in both ventral and lateral regions, meaning that cells spent an average of 49 min across a 2 hour timelapse in a T1 transition (Fig 2F). It is possible for cells to go through a higher number of transitions at a faster rate, or a smaller number of transitions at a slower rate, and spend the same fraction of time in a T1 transition state. Therefore, we tested whether there were differences in rapid junction remodeling versus prolonged rearrangement states by measuring the average time cells remained in a single T1 event before completing a transition. Cells exhibited similar T1 durations in both ventral and lateral regions, with an average T1 duration of 71 minutes for the first transition (78 minutes in lateral regions and 64 minutes in ventral regions) (Fig 2G; Fig S2A-B). For each consecutive transition, the average duration of each T1 transition decreased, with a mean dwell time of 19 minutes for cells completing a fourth transition (23 minutes in lateral regions and 15 minutes in ventral regions) (Fig 2G; Fig S2A-B).

We quantified cell shape change over time, focusing on aspect ratio as a measure of anisotropic tension (Mongera et al., 2018). Ventral cell aspect ratios increased over time with an average slope of 0.0016 min^-1^ across an hour and a half, indicating a gradual but sustained elongation of ventral cells (Fig 2H; See Video S5). Cell elongation parallels the strain pattern observed in brightfield images, where ventral cells undergo axial compression, which may facilitate passive neighbor exchanges (Anjum et al., 2025). By contrast to ventral cells, cells in lateral regions retain a more polygonal shape, indicative of a more isotropic tensions (Fig 2D)(Zitong et al., 2024). The lack of significant differences in T1 transitions between these regions suggests that cell rearrangements are passive responses rather than actively driving rotational and compression in lateral and ventral regions. These results are supportive of counter-rotational vortices arising from external forces and boundary constraints. To further investigate the nature of such a boundary or how forces are distributed within these tissues, we examined the cytoskeletal architecture and extracellular matrix organization, both of which are known to regulate tissue stiffness, adhesion, and large-scale morphogenetic movements (Shawky and Davidson, 2015).

### Emergence of the Supracellular Cytoskeletal and ECM Structures

Immunostaining revealed a complex meshwork of cytoskeleton and ECM in posterior tissues (Fig 3A). Keratin was absent from the blastopore and dorsal midline, while actin fibers were concentrated in dorsal neural cells and along the margins of the ventral blastopore (Fig 3A). Notably, we did not observe a purse-string-like actin structure, a feature surrounding the posterior in rotating fly genitalia (Sato et al., 2015), or proposed as the driving force for counter-rotating vortices during chick gastrulation (Fig 3A)(Xiong et al., 2020). The lack of overtly organized F-actin suggests that counter-rotational movements in *Xenopus* embryos arises through mechanisms distinct from those observed in avian gastrulation. Keratin organization increased in density and bundling as development progressed, making individual fibrils increasingly difficult to resolve by the tailbud stages (stage 24; Fig 3B-C). Additionally, apically localized keratin fibers in ventral epithelial cells aligned perpendicular to the mediolateral axis (Fig 3B-C).

**Figure 3.**
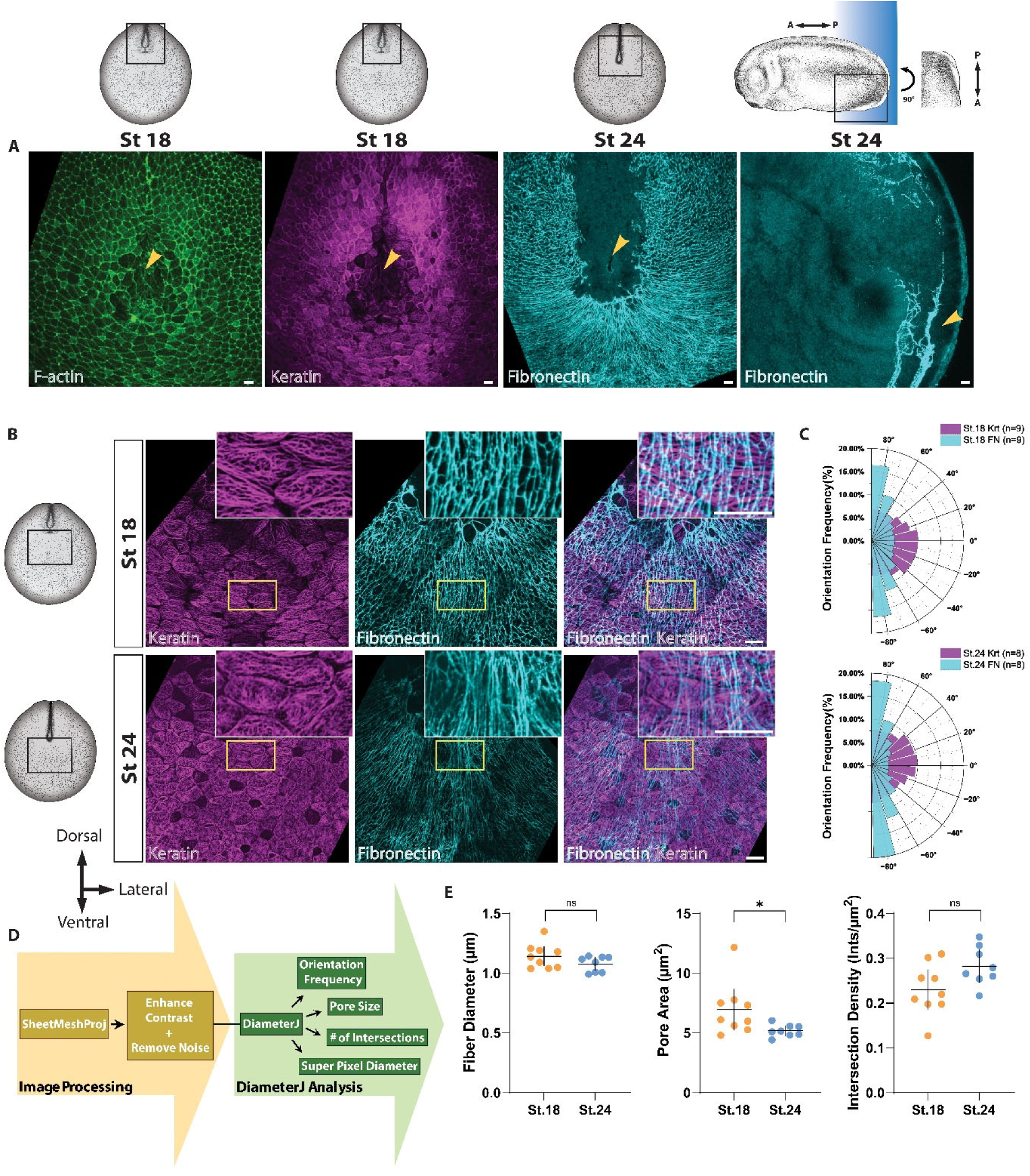
Cytoskeletal and extracellular matrix remodeling during posterior development. (A-B) 2D-projected confocal images of the near-blastopore region (orange arrowhead) at stages 18 and 24. Schematics indicates approximate imaging locations and plane (Blue shaded box). (A) En face views of F-actin, keratin 8, and fibronectin and fibronectin in parasagittal section. The sagittal section is rotated 90° to align the blastopore with en face images. The regions of tissue rotation are located approximately 100 to 150 µm from the center of the blastopore. (B) En face views of keratin 8 and fibronectin, with a zoomed inset (yellow box). Aligned fibronectin fibrils are most prominent within 80 to 120 µm of the blastopore. (C) Polar histograms (rose plots) of fiber orientations at stage 18 and 24 (n, number of embryos used; Square goodness-of-fit test, ∗∗∗∗p<0.0001) (D) Workflow for image processing and analysis of immunostained samples. (E) Fibronectin morphological features from stage 18 and stage 24. (C and E) measured within a 100 by 100 µm region ventral to the blastopore. Each symbol represents the mean value per embryo (Mann-Whitney U, ∗p=0.02). Bars; mean ± 95% CI. (A-B) scale bars, 20 µm. (A-B) Xenopus illustrations © Natalya Zahn (2022)(Zahn et al., 2022).

Given the role of the ECM in providing structural support and integrating mechanical forces across tissues (Palmquist et al., 2022), we next examined its composition and distribution. Collagen and fibrillin, key ECM components involved in structural organization and tissue mechanics (Skoglund, 1996), were observed surrounding the notochord and presomitic mesoderm but were notably absent from the most posterior regions near the blastopore between stages 18 to 22 (Fig S3A). Thus, collagen and fibrillin do not play a role in regions near the blastopore during early tailbud stages. Fibronectin was mostly absent from neural tissues and dorsal to the blastopore but formed a dense fibrillar network approximately 100 µm ventral of the blastopore. In these ventral regions, the fibronectin network assumed a fan-like morphology with long straight fibers extended radially away from the blastopore, with thinner collateral fibers woven through and spanning gaps between long fibers (Fig 3A). As observed in both earlier (Davidson et al., 2004) and later developmental stages (Jackson et al., 2017), fibronectin fibrils in posterior tissues organize into two distinct layers (Fig 3A). A dense fibrillar layer forms at the ectoderm-mesoderm interface (e.g., superficial-most), while a looser and more dispersed network develops between the mesoderm and endoderm (Fig S4A-B). Fibers in the superficial-most layer appear to connect with fibers in the deeper layer within the blastopore collar (Fig 3A). Fibronectin was also found to closely associate with laminin in ventral posterior regions (Fig S3B).

To distinguish morphological changes in the superficial fibronectin layer from deeper networks, we computationally isolated the superficial layer and analyzed it separately (Fig 3D; Fig S4A-C). Statistical analysis of fiber orientation showed a stronger orientation preference over time with increased clustering of fibers around the mean direction at stage 24 (R = 0.38) compared to Stage 18 (R = 0.28) (Fig 3B-C). This was further supported by an increase in the concentration parameter (κ = 0.92 at stage 24 vs. κ = 0.67 at stage 18), reflecting a progressive alignment of fibronectin fibrils as the tailbud develops (Fig 3B-C). We also noted that fibronectin fibril alignment was perpendicular to keratin fiber alignment located in the superficial epithelium, two cell layers above (Fig 3B-C). Further analysis of fibronectin fibril architecture revealed 0.05 intersections/µm increase in fiber intersection density and a 0.03 µm decrease in fiber diameter by stage 24 (Fig 3B,E). The most significant structural change was a 0.97 µm² reduction in pore size (Fig 3B,E), which could be due to increased interconnectivity. These findings indicate that from stage 18 to stage 24, the fibronectin network undergoes structural remodeling, becoming both denser and more directionally aligned.

### Counter-Rotations Depend on Dorsal-Ventral patterning

The occurrence of aligned fibrils to ventral tissues despite broad *fn1* expression (Briggs et al., 2018), raises the possibility that dorsal and ventral regions may differ in how they assembly with or remodel the extracellular matrix. To investigate this further, we generated dorsalized and ventralized embryos by treating with LiCl or UV, respectively, each distinctly lacking either dorsal or ventral structures (Kao and Elinson, 1989; Shook et al., 2022) (Fig 4A-B; See STAR Methods). Coordinated flow patterns in both dorsalized and ventralized were disrupted with a loss of counter-rotational flows (Fig 4C; See Video S6). In ventralized embryos, vorticity appeared reduced and largely homogeneous across the tissue (Fig 4D). By contrast, dorsalized embryos exhibited small, localized regions of opposing vorticity, lacking any large-scale, structured counter-rotation (Fig 4D). Consistent with the loss of organized counter-rotational flows, SWIRL analysis detected only 8 total vortices in 11 dorsalized embryos and 11 vortices in 11 ventralized embryos, compared to 32 vortices in 19 wild-type embryos (Fig 4E). Vortices in dorsalized and ventralized embryos were less compact, rising from an average of 0.08 in control embryos to 0.34 in ventralized embryos, and 0.23 in dorsalized embryos (Fig 4F).

**Figure 4.**
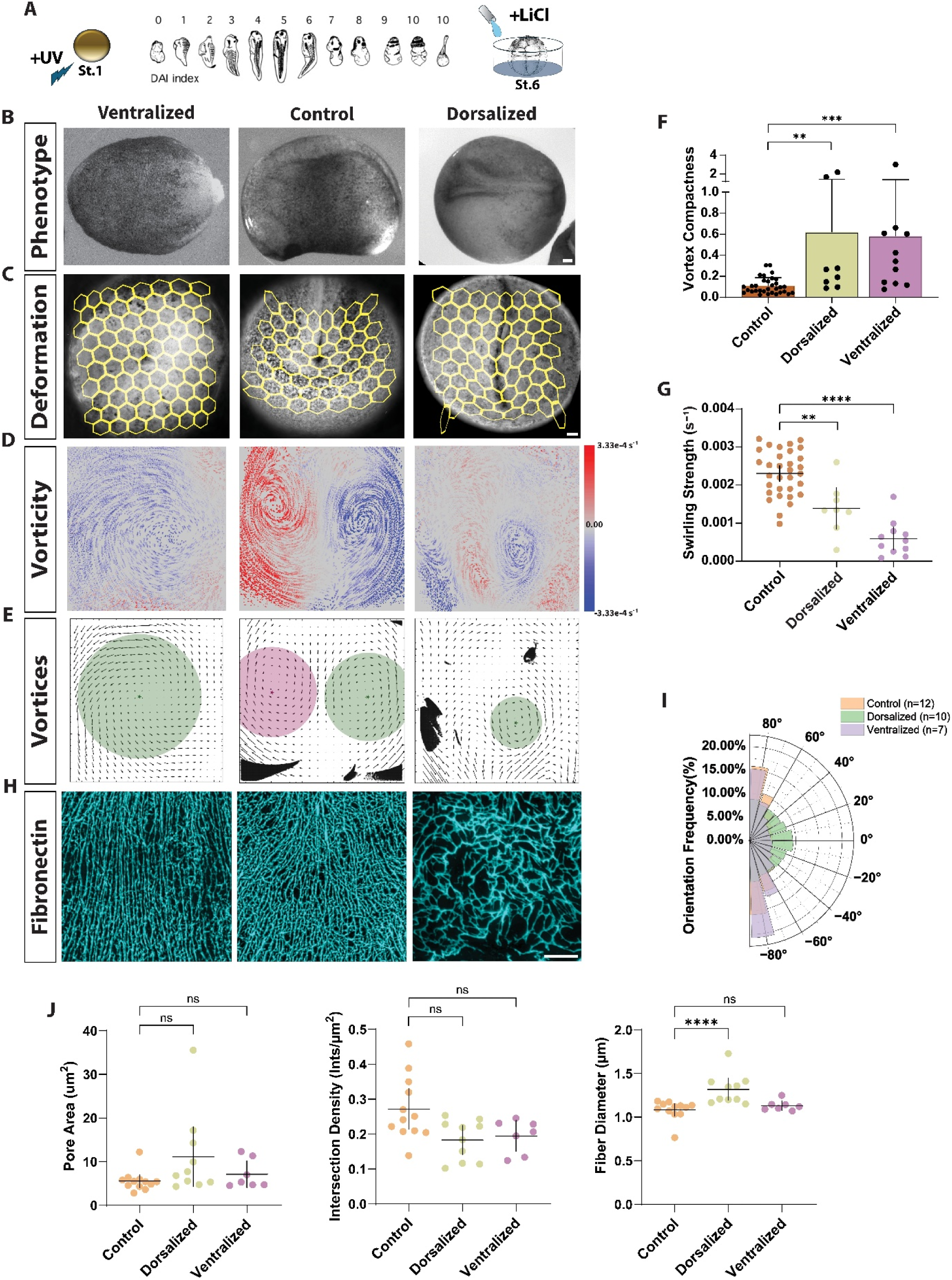
Ventralized and dorsalized tissues exhibit distinct flow dynamics and fibronectin organization. (A) Dorso-anterior index (DAI) of the severity of dorsalized and ventralized phenotypes(Kao and Elinson, 1988). (B) Representative phenotypes of each treatment at stage 20, showing characteristic morphological differences between dorsalized, ventralized, and control embryos. (C) Final frame of brightfield timelapse sequence, overlaid with a yellow deformation map (see also Video S6). (D) Vorticity overlaid randomized dot plots deformed by calculated displacements, and max-projected across all timepoints to visualize movement patterns. Representative images are shown for each treatment. (E) SWIRL predicted vortices highlight vortex structure and organization. (F-G) Quantitative comparison of predicted vortex characteristics in dorsalized (8 vortices from 11 embryos), control (32 vortices from 19 embryos), and ventralized (11 vortices from 11 embryos) embryos. Each symbol represents a single predicted vortex (Mann-Whitney U, ∗∗p=0.0012; ∗∗p=0.0061; ∗∗∗p=0.0002; ∗∗∗∗p<0.0001). (F) Vortex compactness (mean ± 95% CI). (G) Vortex swirling strength (mean ± 95% CI). (H) Fibronectin networks in posterior tissues from stage 24 embryos within a 100 by 100 µm region ventral to the blastopore. (I) Fibronectin orientation frequency in each treatment (n, number of embryos used; Square goodness-of-fit test, ∗∗∗∗p<0.0001) (J) Morphological features of fibronectin network in dorsalized, control, and ventralized embryos. Each symbol represents the mean value per embryo (Mann-Whitney U, ∗p=0.0426; ∗∗∗∗ p < 0.0001). Bars indicate mean ± 95% CI. (B-C) scale bars, 100 µm; 20 µm in (H). (A) DAI diagram with permission of the publisher. Xenopus illustrations © Natalya Zahn (2022)(Zahn et al., 2022).

Swirling strength was also reduced in both treatments, decreasing from 0.0023 s⁻¹ in control embryos to 0.0013 s⁻¹ in dorsalized tissues, and 0.0005 s⁻¹ in ventralized tissues (Fig 4G). Changes in vortex compactness and strength indicate a loss of the well-structured rotational motion in both ventralized and dorsalized embryos (Fig 4E-G). Taken together, alterations in dorsal or ventral identities point out the role of large scale DV patterning in maintaining organized counter-rotational flow.

Given the possible role of fibronectin in coordinating cell movements, we next analyzed fibronectin organization in dorsalized and ventralized embryos (See STAR Methods). Fibronectin networks were highly dense and well-aligned in ventralized tissues, particularly surrounding the blastopore (Fig 4H). By contrast, fibronectin in dorsalized embryos was sparse and disorganized, resembling fibronectin networks found in lateral tissues adjacent to neural structures in wild-type embryos (Fig 4H). Orientation analysis reveals that fibronectin fibrils in ventralized embryos were more aligned than in control samples, whereas fibronectin fibrils in dorsalized embryos showed no clear alignment (Fig 4I). The diameter of fibronectin fibrils increased by approximately 0.116 µm in dorsalized tissues from controls (Fig 4J). Interestingly, the number of fiber intersections decreases by 0.044 intersections per µm^2^ in dorsalized embryos, and 0.039 intersections per µm^2^ in ventralized samples, suggesting that connections in fibronectin are maintained, but the fibers in dorsalized embryos are not fully co-aligned or actively cross-linked (Fig 4J). Fibronectin pore sizes in both control and ventralized embryos appeared smaller, yet these pores were elongated and narrow, with few collateral fibers bridging the radial fibers (Fig 4H,J). The complete loss of counter-rotations in both dorsalized and ventralized embryos strongly suggests that the interaction between dorsal and ventral cell identities is essential for the emergence of these flows. Individual cells in the superficial ectoderm undergo transitions that may contribute to directed force generation, as well as increase local tissue density, potentially influencing tissue stiffness (Asai et al., 2024).

### Counter-Rotations Persist Despite Cell Behavior Perturbations

To investigate the role of cell division, radial intercalation, and convergent extension in tissue mechanics and counter-rotational flows, we implemented pharmacological inhibitors, as well as targeted genetic perturbations (Fig 5A-B; See STAR Methods). We first blocked cell division with a combination of hydroxyurea and aphidicolin (HUA), which arrests cell cycle progression through the S phase by inhibiting DNA synthesis (Pokrovsky et al., 2021). This treatment resulted in larger cell sizes, consistent with a compensatory increase in cell volume due to reduced proliferation (Fig S5A)(Mitsui and Schneider, 1976). Despite this reduction in overall cell density, embryos exhibited no major morphological defects at neural and tailbud stages, consistent with prior research (Fig 5C)(Harris and Hartenstein, 1991).

**Figure 5.**
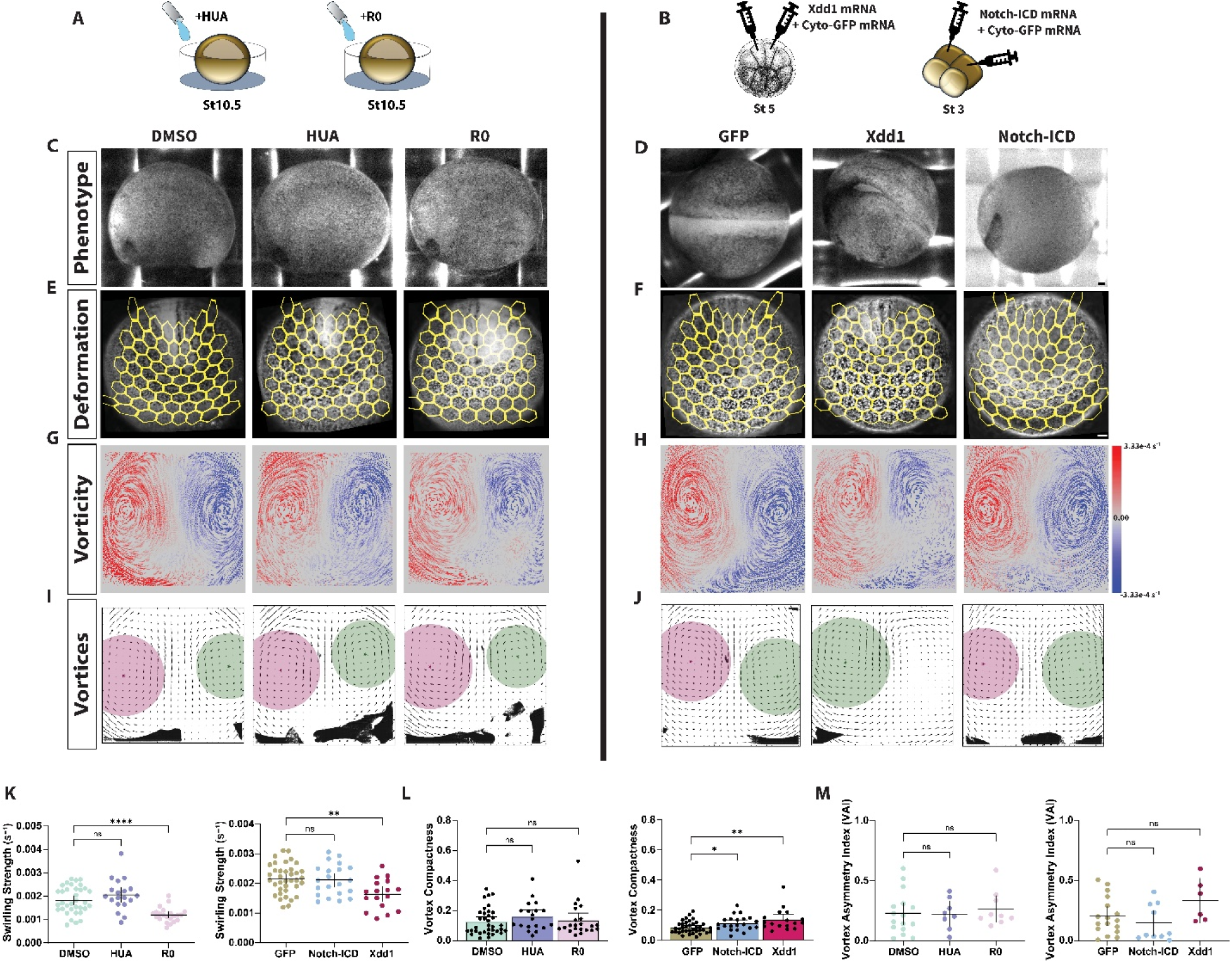
Disrupting cell division, convergent extension, and radial intercalation has minimal effects on vortex formation. (A-B) Methods used to perturb specific cellular behaviors in dorsal and ventral tissues. (C-D) Representative phenotypes showing minimal morphological differences across conditions. (E-F) Final frame of timelapse sequence overlaid with a yellow deformation map indicating cumulative deformation. (G-H) Time-projected view of vorticity overlaid onto randomized dot plots deformed using calculated displacements. Across treatments, vorticity patterns remained largely unchanged, consistent with the retention of counter-rotational flows. (I-J) SWIRL predicted vortices after treatment. (K-M) Vortex characteristics across treatments. In (K) and (L), each symbol represents a single predicted vortex: DMSO (35 vortices from 19 embryos), HUA (19 vortices from 11 embryos), R0 (22 vortices from 13 embryos), GFP (37 vortices from 20 embryos), Xdd1 (17 vortices from 11 embryos), and Notch-ICD (21 vortices from 11 embryos). In (M), each symbol represents an embryo with a vortex pair (Mann-Whitney U, ∗p=0.0247; ∗∗p=0.0012; ∗∗p=0.0016; ∗∗∗∗p<0.0001). Bars indicate mean ± 95% CI, except in (L). (C-D) scale bar, 100 µm. (A-B) Xenopus illustrations © Natalya Zahn (2022)(Zahn et al., 2022).

Radial intercalation of multiciliated cells significantly influences tissue architecture by driving cell movements from deeper layers toward the epithelial surface, thereby altering tissue density and mechanics. We manipulated radial intercalation via targeted modulation of Notch signaling with RO4929097(R0), a γ-secretase inhibitor, to enhance radial intercalation by increasing the number of differentiating multiciliated cells (Myers et al., 2014)(Fig 5A; See STAR Methods). RO4929097 blocks the proteolytic cleavage of the Notch receptor, preventing release of the Notch intracellular domain (Notch-ICD) and subsequent activation of downstream target genes. This disruption in Notch signaling relieves lateral inhibition, promoting excessive differentiation and apical migration of multiciliated cells. To suppress radial intercalation we employed Notch-ICD overexpression (See STAR Methods). Notch-ICD specifically inhibits differentiation and intercalation of multiciliated cells (Chitnis et al., 1995), thereby reducing cell movement and altering epithelial composition and density. To isolate changes in tissue density while minimizing effects on other tissues, we targeted Notch-ICD mRNA to the ventral ectoderm (Fig 5B). RO-treated embryos showed a clear increase in surface-localized multiciliated cells (Fig S5B-C). By contrast, Notch-ICD–injected embryos exhibited a marked depletion of surface ciliated cells (Fig S5D). Despite the expected alterations in cell intercalations, neither treatment resulted in significant morphological disruptions (Fig 5C-D).

Finally, to examine the role of convergent extension in driving posterior movements, we disrupted the planar cell polarity (PCP) pathway by targeting *Xdd1* mRNA to the neural plate (see STAR Methods). *Xdd1* acts as a dominant-negative form of Dishevelled, a core PCP component, which consequently blocks downstream PCP signaling required for mediolateral intercalation (Sokol, 1996). As expected, *Xdd1* overexpression blocked medio-lateral intercalations, preventing neural tube fusion and convergent extension (Fig 5D).

Despite reducing cell division, altering the number of cells intercalating radially into the epithelium, and disrupting convergent extension processes, overall patterns of tissue deformation and posterior morphogenesis remained largely unchanged (Fig 5E-F). Across all perturbations, embryos retained the bi-directional vorticity pattern, indicating that counter-rotational flows persist even when specific cell behaviors are disrupted (Fig 5G-H). However, vortex analysis revealed subtle changes in vortex properties (Fig 5I-M). Specifically, swirling strength reduces from an average of 0.0017 s^-1^ in DMSO controls, to 0.0011 s^-1^ in embryos with a global increase of ciliated cells (R0) (Fig 5K). Similarly, vortex compactness increased to 0.11 in embryos with reduced ventral radial intercalations (Notch-ICD), indicating that vortices were less tightly organized (Fig 5L). To distinguish changes in the overall symmetry of vortices, we also compared the swirling strength of vortices found in the same embryo. We observed no changes in vortex asymmetry (Vortex Asymmetry Index; VAI; see Supplemental Material S1) from these treatments, ranging from a median VAI of 0.05 in Notch-ICD injected embryos, 0.17 in GFP controls, 0.21 in both R0 and HUA treated embryos, and DMSO controls at 0.18 (Fig 5M). These results further implicate tissue density and stiffness contribute to the maintenance of coordinated counter-rotations, particularly in relation to the boundary conditions established in ventral tissues.

We observed a similar decrease in vorticity strength and symmetry in embryos with disrupted convergent extension (*Xdd1*). Specifically, swirling strength decreased from 0.0021 s⁻¹ in GFP controls to 0.0017 s⁻¹ in *Xdd1* embryos (Fig 5K). These results suggest that the loss of mediolateral intercalation weakens the posteriorly directed movements required to generate counter-rotations. This effect was accompanied by increased vortex compactness, with an increase from 0.079 in GFP controls to 0.116, suggesting that dorsal convergent extension contributes not only to force generation but also contribute to the spatial organization of rotational tissue flows (Fig 5L). We did observe a slight increase in the asymmetry of vortices in *Xdd1* injected embryos, from 0.16 in GFP controls, to 0.30, however, this difference was not significant (Fig 5M).

### Ventral Fibronectin Regulates Counter-Rotational Flows

To test the role of fibronectin in posterior morphogenesis by disrupting its structural role or its ability to interact with cells, we used two approaches. First, we used the integrin β1 function-blocking antibody P8D4, which prevents integrin-fibronectin binding, leaving the fibronectin network intact but removing its mechanical linkage to cells (Davidson et al., 2006). To specifically target the posterior fibronectin–cell connections at neurula stages, P8D4 was microinjected into the archenteron during late gastrulation stages, allowing diffusion into the posterior region before epithelial and mesodermal rearrangements become fully established (See STAR Methods). Second, we targeted fibronectin antisense morpholinos (FNMO) injections to the ventral ectoderm to knock down fibronectin expression (Davidson et al., 2006), reducing fibril deposition, while preserving fibronectin and integrin-mediated attachment in dorsal tissues (Fig 6A; Fig S6B; See STAR Methods). By late neural and early tailbud stages, P8D4- and FNMO-treated embryos developed similar defects, with P8D4-injected embryos displaying more severe phenotypes (Fig 6B). In both conditions, ventral ectoderm exhibited localized tissue bunching, suggesting a loss of substrate for mesoderm migration (Fig 6B) (Davidson et al., 2004). In FNMO-injected embryos, the dorsal ectoderm appears to be normal, whereas in P8D4-injected embryos, the dorsal ectoderm detached from the ventral mesoderm (Fig 6B). The greater severity of defects in P8D4-treated embryos is likely due to the broader disruption of integrin-mediated adhesion, which affects a larger number of cells and compromises the mechanical coupling between fibronectin and surrounding tissues.

**Figure 6.**
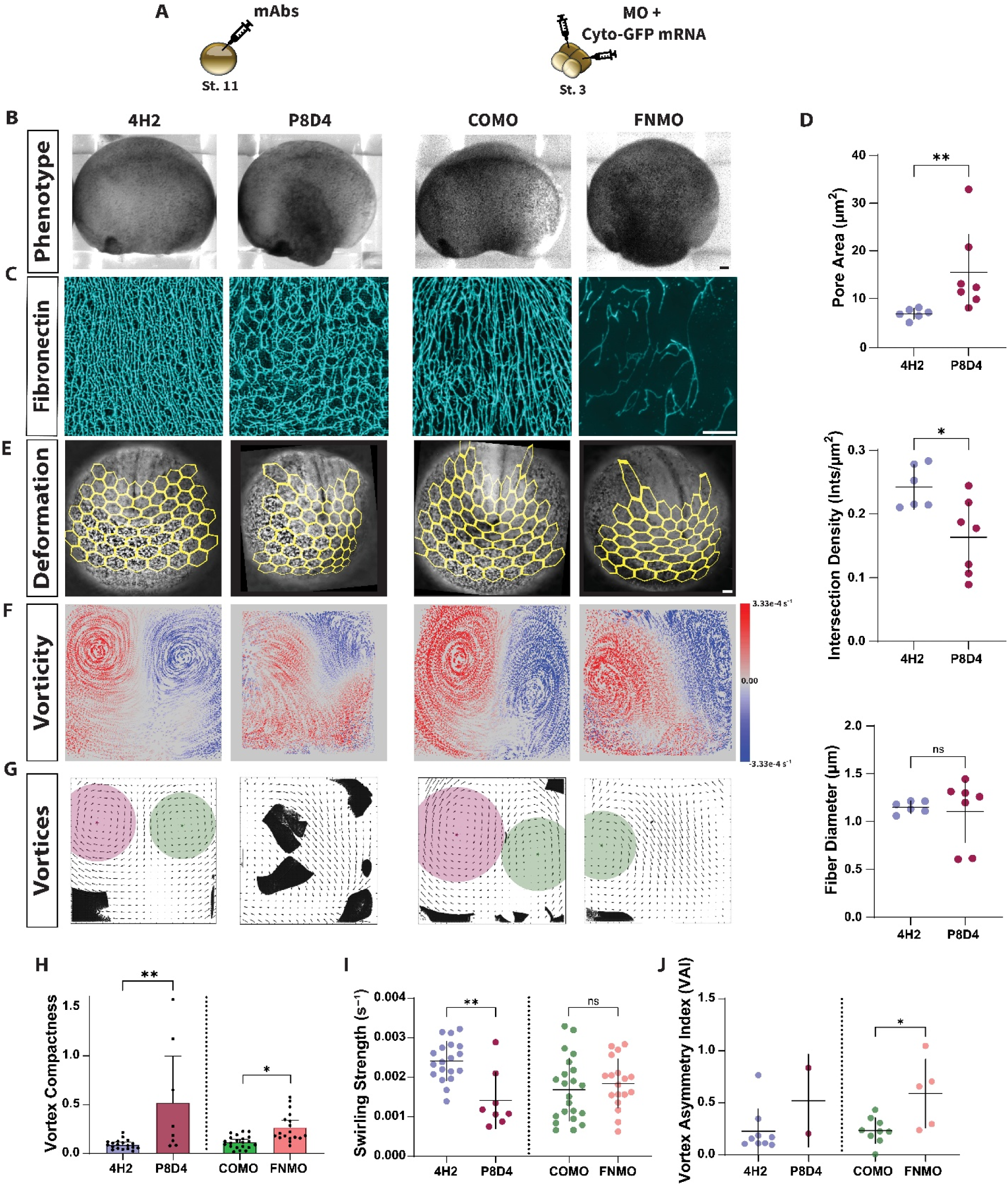
P8D4 and Morpholino treatments disrupt vortex formation and fibronectin organization. (A) Methods used for targeted disruption of fibronectin organization and integrin-fibronectin interactions. (B) Representative phenotypes showing morphological changes following fibronectin disruption. (C) 2D max-projected confocal image of fibronectin within a 100 by 100 µm region ventral to the blastopore. (D) Morphological features of fibronectin matrix for control (mAb 4H2) and function-blocking (mAb P8D4) treatments. Each symbol represents the per-embryo mean (Mann-Whitney U, ∗p=0.0350; ∗∗p=0.0023). Bars indicate mean ± 95% CI. (E) Final frame of brightfield timelapse sequence overlaid with yellow deformation map (see also Video S7). (F) Time-projected displacement of random dot plot overlaid with vorticity. Disruptions to fibronectin result in less distinct or absent bi-directional vortices compared to controls. (G) SWIRL predicted vortex structure and spatial distribution across treatments. (H-J) Vortex characteristics across treatments. Each symbol in (H) and (I) represents a single predicted vortex: 4H2 (19 vortices from 10 embryos), P8D4 (8 from 11), COMO (21 from 13), and FNMO (18 from 13). In (J), each symbol represents an embryo with a predicted vortex pair. Vortex formation was significantly disrupted in P8D4- and FNMO-treated embryos, with a reduction in detected vortices and alterations in vortex compactness (H) and swirling strength (I). (J) Vortex asymmetry index (Mann-Whitney U, ∗p<0.0290; ∗∗p=0.0054; ∗∗p=0.0024; ∗∗∗p=0.0002). Bars indicate mean ± 95% CI, except in (H). Scale bars, 100 µm in (B,E); 20 µm in (C).

Immunostaining confirmed significant disruptions to fibronectin organization following both treatments. FNMO-injected embryos exhibited a clear reduction in fibronectin fibril formation, with only sparse fibrils present in targeted ventral tissues (Fig 6C; Fig S6A). By contrast, P8D4-treated embryos displayed more severe disruptions, where fibronectin fibrils were present, but failed to form the highly organized networks observed in control embryos (Fig 6C; Fig S6B). While fiber diameter remained unchanged, pore sizes increased by 5.4µm in P8D4-treated embryos, and the density of fiber intersections was reduced by 0.05 intersections per µm^2^ (Fig 6D). Blocking integrin interaction reduced connectivity between fibrils and increases the spacing between fibrils, suggesting that the matrix may not have evolved beyond the time P8D4 was injected (Fig 6C). Additionally, sagittal sections revealed the detachment of deep ventral cells in P8D4-treated embryos (Fig S6B); delamination was not observed in FNMO embryos.

Both P8D4-mediated disruption of integrin-fibronectin interactions and FNMO-mediated reduction of fibronectin fibril formation significantly affected posterior tissue deformation and rotational flow patterns (Fig 6E; See Video S7). In both conditions, counter-rotational movements were disrupted, although the severity of disruption varied. In P8D4-treated embryos, vortex separation was nearly eliminated, leading to a complete loss of distinct rotational flows (Fig 6F). By contrast, FNMO-injected embryos retained a weaker separation of rotational flows, but vortex stability and coherence were impaired (Fig 6F). Quantitative vortex analysis revealed a reduction in the number of vortices detected with only 8 vortices detected across 11 P8D4-treated embryos, compared to 19 vortices across 10 control embryos (Fig 6G). The remaining vortices exhibited increased compactness (0.09 to 0.23) (Fig 6H) and decreased swirling strength (0.0023 s⁻¹ to 0.0011 s⁻¹) (Fig 6I). These results indicate that integrin-fibronectin anchoring is necessary for maintaining distinct and structured counter-rotational movements. FNMO-treated embryos, on the other hand, showed only a slight reduction in vortex number (18 vortices across 13 FNMO-treated embryos, compared to 21 vortices across 13 COMO embryos) (Fig 6G). However, like P8D4-treated embryos, generated vortices were less structured with compactness values increasing from 0.11 in controls, to 0.19 (Fig 6H). Notably, swirling strength was not reduced in FNMO-treated embryos, however, an increase in VAI (0.23 to 0.66) indicates an imbalance in rotational flow dynamics (Fig 6I-J). These results indicate that direct integrin-fibronectin interactions are necessary to anchor ventral tissues together and provide mechanical support during large-scale posterior movements.

## Discussion

We have identified counter-rotational flow in the posterior *Xenopus* embryo during tailbud morphogenesis and have tested a set of cell behaviors that have been suggested for driving such flows in other systems. Using live imaging, quantitative deformation analysis, and computational vortex detection, we characterized the emergence and persistence of rotational flows. Perturbation of cell behaviors and fibronectin suggest that the physical constraint of ventral ECM plays a key mechanical role in posterior tailbud morphogenesis. Our study further suggests counter-rotational flows depend on patterning regions with fluid- and solid-like mechanics and offers insights into how supracellular mechanics shape vertebrate development.

Our analysis of nuclear velocities suggests that rotational flows are consistent with an externally-driven forced vortex. Disruption of cell division, radial intercalation, and convergent extension revealed that these cellular behaviors influence vortex organization but are not primary drivers of counter-rotational flow. While inhibition of cell division and intercalation altered vortex properties, such as reducing swirling strength or increasing compactness, the overall presence of counter-rotational flow remained unchanged (Fig 5E-M). Similarly, disruption of planar cell polarity signaling via Xdd1, which impairs mediolateral intercalation in the dorsal neural plate, did not abolish rotational flow. Although these embryos exhibited defects in neural tube closure, posterior axis elongation still occurred, consistent with previous findings that trunk convergent extension and posterior elongation are decoupled (Wallingford and Harland, 2002; Tao et al., 2005). Other compensatory mechanisms may be at play, helping to maintain tissue-scale mechanical properties even when specific cellular processes are perturbed. These findings reinforce the view that rotational movement is mechanically regulated through long range tissue-scale forces.

Radially oriented fibronectin and other ECM ventral to the blastopore are needed for distant counter-rotational movements (Fig 3A) and may provide structural support for the blastopore for its translocation to the ventral side of the embryo. While not as aligned, fibronectin and other ECM play a major role in somitogenesis on the dorsal side (Guillon et al., 2020). Increased fibronectin density promotes the clustering of α5β1 integrins, which strengthens adhesion and enables cells to generate and sustain traction forces (Roca-Cusachs et al., 2009). Highly organized fibronectin likely enhances cell adhesion, increases tissue stiffness, and reinforces ventral structural integrity (Singh et al., 2010). Additionally, fibronectin, laminin, and other ECM can act as force transmission elements, facilitating mechanotransduction and enabling cells to sense and respond to their mechanical environment (Scott et al., 2015). The role of fibronectin in shaping posterior morphogenesis is underscored by our ECM perturbation experiments. Blocking integrin-fibronectin interactions led to a near-total disruption of counter-rotational flows, while fibronectin knockdown resulted in a reduction in vortex organization (Fig 6D-F,H). These findings suggest that the observed fibronectin architecture in ventral tissues may not simply reflect passive mechanical changes but could actively shape the forces driving rotational movements by modulating adhesion strength, tissue stiffness, and mechanotransduction pathways.

Based on our observations and experimental results, we propose a two-component model for the emergence of counter-rotational flows. First, posteriorly directed movements from convergent extension drive large-scale tissue displacement, which, in isolation, is insufficient to generate counter-rotational flow. Instead, counter-rotations emerge as these posteriorly directed movements interact with a mechanical boundary. This boundary may be established through fibronectin-mediated adhesion and structural support or by local increases in tissue density driven by cell behaviors such as cell division or radial intercalation. Fibronectin could serve as a scaffold that organizes tissue interactions and constrains movement patterns, while cellular rearrangements may generate local resistance to posterior tissue flows, reinforcing the boundary condition necessary for counter-rotations to arise.

This mechanical framework aligns with a critical developmental transition, as counter-rotational flows emerge just as anterior morphogenesis concludes and tail formation begins (Masak and Davidson, 2023). Neural tube closure marks the end of primary neurulation and positions neural mesodermal progenitors (NMPs) between the posterior end of the neural tube and the dorsal lip of the blastopore (Gont et al., 1993). Following neural tube closure, somitogenesis (Miao and Pourquie, 2024) coordinates with secondary neurulation processes (Lowery and Sive, 2004; Catala, 2021), utilizing these progenitors to generate the tail. This spatial reorganization coincides with the emergence of large-scale rotational flows, ventral tissue compression, and fibronectin alignment, and together, these features suggest a broader shift in the mechanical landscape of the posterior embryo. Therefore, these morphogenetic forces likely shape not only tissue geometry, but also the local environment in which NMPs are maintained, positioned, and integrated into the elongating axis.

In summary, we have identified a novel counter-rotational flow pattern in the posterior *Xenopus* embryo that emerges through externally driven mechanical forces rather than intrinsic cellular organization. Our results demonstrate that dorsal-ventral patterning, ECM interactions, and large-scale tissue mechanics collectively establish the conditions necessary for these flows. By integrating concepts from developmental biology and fluid dynamics, this study advances our understanding of how global morphogenetic movements shape posterior body formation, with potential implications for developmental biology and tissue engineering.

## Acknowledgements

We thank members of the Davidson laboratory for their support and feedback throughout this work. In particular, we are grateful to Jing, Carsten, and Sommer for their critical discussions, technical assistance, and thoughtful insights at every stage of this project. We also thank the Developmental Studies Hybridoma Bank (DSHB) for their guidance on antibody purification and concentration. This work was supported by grants to LAD from the Eunice Kennedy Shriver National Institute of Child Health and Human Development at the National Institutes of Health (R37 HD044750 and R21 HD106629).

## Data and code availability

All data reported in this paper will be shared by the lead contact upon request. All original code has been deposited at Harvard Dataverse and is publicly available at DOI:10.7910/DVN/5NBKQU as of the date of publication. Any additional information required to reanalyze the data reported in this paper is available from the lead contact upon request.

## Materials and Methods

### *Xenopus laevis* embryos and mRNA microinjections

All studies conducted using *Xenopus laevis* embryos were approved by the University of Pittsburgh Division of Laboratory Animal Resources. *Xenopus* embryos were collected, in vitro fertilized, de-jellied, and microinjected with mRNAs for fluorescent probes using methods described previously (Newman et al., 2018). Transgenic *CMV:memGFP* frogs were obtained from the National *Xenopus* Resource. Embryos were staged according to Nieuwkoop and Faber (NF)(Nieuwkoop PD, 1956). All frog animal use protocols were in compliance with Public Health Service (PHS) and United States Department of Agriculture (USDA) guidelines for laboratory animal welfare and reviewed and approved by the University of Pittsburgh Institutional Animal Care and Use Committee (IACUC).

For microinjection of mRNAs, embryos were placed in 1X Modified Barth’s Solution (MBS) with 0.3% Ficoll. To visualize nuclei, embryos were injected at the single-cell stage with 200 pg of H2B-mScarlet mRNA. To track cells expressing dominant-negative constructs and morpholinos, 60 pg of 3X-EGFP mRNA was co-injected as a tracer. To block convergent extension, up to 1ng of Xdd1 mRNA was injected at the 16-cell stage, targeted in dorsal medial animal blastomeres (Wallingford and Harland, 2002). To block cell intercalation in ventral tissues, 34 pg of Notch-ICD mRNA was injected into ventral animal blastomeres at the 4-cell stage (Chitnis et al., 1995). By stage 17, embryos were sorted based on the expected fluorescence expression pattern. Effective disruption of dorsal extension was confirmed by the resulting embryo phenotypes, with neural tube closure defects consistent with previously reported findings (Wallingford and Harland, 2002). To assess the effectiveness of Notch-ICD in blocking cell intercalation, ciliated cells were visualized and quantified via acetylated-tubulin immunostaining (see Supplemental Methods).

### Knockdown / Morpholinos

Antisense morpholinos against fibronectin were synthesized by GeneTools, LLC (Philomath, OR) as previously described (Davidson et al., 2006).The sequences are as follows: XFN1.L MO(5’-CGCTCTGGAGACTATAAAAGCCAAT-3’); XFN1.S MO (5’-CGCATTTTTCAAACGCTCTGAAGAC-3’). The XFN1.L and XFN1.S morpholinos were combined (50:50, from 1 mM stock solutions) in water and injected at up to 4.8µmol of each morpholino, or 9.6 µmol total per embryo. 60pg of 3X-GFP mRNA was co-injected as a tracer. We used the standard control morpholino (Gene Tools) 5’-CCTCTTACCTCAGTTACAATTTATA-3’. We targeted the posterior ventral by injecting ventral animal blastomeres at the 4-cell stage (Davidson et al., 2006). Knockdown effectiveness was assessed by fibril morphology (Fig S6A).

### Embryos embedded in agarose for imaging

An acrylic chamber is prepared with a glass coverslip, using vacuum grease to adhere the glass and create a seal at the bottom of the chamber. Ultra-low melting temperature agarose 1.5% w/v is prepared in 1/3X MBS and heated to 65°C. The solution is pipetted into the prepared chamber and cooled down to 30°C–37°C, before stage 18 embryos are transferred into the chamber. Hair tools are used to position embryos in the agarose as it continues to cool so that the blastopore is positioned adjacent to the bottom of the chamber (Fig 1B). 1/3X MBS is added to fill the chamber once the agarose has fully cooled, then sealed with a coverslip (Chu and Davidson, 2022).

### Purification, concentration and microinjection of function-blocking antibodies

To disrupt integrin binding to fibronectin, 10ml of function-blocking mAb P8D4 (19ug/µl) and control mAb 4H2 (18ug/µl) were obtained from Developmental Studies Hybridoma Bank (DSHB) and were purified using 30kDa MWCO Cytiva Protein G HP SpinTrap™ Columns (28903134; Cytiva) and concentrated using Amicon® Ultra-4 filter devices (UFC8030; Sigma), following the manufacturer’s protocol for each. Final concentrations of 4H2 and P8D4 were measured by a Nanodrop device, estimated to be 0.6 mg/ml and 1.3 mg/ml, respectively. Embryos were injected with up to 80ng of antibodies into the blastocoel at gastrulation stages (10.5-12) and cultured to stage 18 for imaging.

### Treatments

Embryos at the start of gastrulation were incubated in 20 mM Hydroxyurea (H8627; Sigma) and 150 μM Aphidicolin (A0781; Sigma), 100 μM RO4929097 (S1575; SelleckChem), or 1% DMSO (vehicle control). Treatment concentrations and culture conditions were selected based on prior *Xenopus* studies (Myers et al., 2014). Hydroxyurea (20 mg/mL) was dissolved in ddH₂O and filtered through a 0.2 μm membrane. Hydroxyurea aliquots were frozen and stored at −20°C in 76 μL aliquots for up to a week. Aphidicolin was reconstituted in DMSO to a stock concentration of 30mM and stored at −20°C in 5 μL aliquots. The filtered Hydroxyurea solution was immediately combined with aliquoted Aphidicolin and diluted in 1% DMSO and 0.1X Marc’s Modified Ringers (MMR) for embryo treatment. RO4929097 was dissolved in DMSO to a stock concentration of 10 mM, stored at −20°C in 20 μL aliquots, and diluted in culture media immediately before use. To confirm cell cycle inhibition, embryos were stained with Phalloidin to label F-actin, verifying an increase in cell size in HUA-treated embryos. The effectiveness of RO4929097 treatment on increasing ciliated-cell density was assessed by α-tubulin labeling following the protocols detailed in Section 3.4.7 and Appendix A3.8. Embryos were dorsalized or ventralized following established protocols (Kao and Elinson, 1989). Only embryos with strong dorsalization (DAI >8) or ventralization (DAI <2) were retained and cultured to stage 18 for time-lapse imaging or stage 24 for immunolabeling.

### Immunofluorescence and Imaging

Embryos were fixed with 4% paraformaldehyde in 1x PBS overnight to visualize F-actin and α-tubulin. For immunofluorescence of ECM (fibronectin, fibrillin, collagen-2, laminin-a1) and keratin, embryos were fixed 5% Trichloroacetic acid (TCA) in 1x PBS overnight at 4°C. Fixed embryos were washed 3X for 30 minutes with PBST (1x PBS + 1% Triton-X) and blocked with 10% CAS-Block (008120; Invitrogen) in PBST for 1 h prior to antibody staining. Immunofluorescence staining was carried out with primary antibodies against fibrillin-2 (JB3, Developmental Studies Hybridoma Bank; 1:200), collagen-2 (II-II6B3, Developmental Studies Hybridoma Bank; 1:100), laminin-a1 (L9393, Sigma-Aldrich; 1:500), mouse anti-fibronectin monoclonal antibody (4H2, courtesy of Douglas DeSimone, University of Virginia, Charlottesville, VA, USA; 1:500), rabbit anti-fibronectin polyclonal antibody (F3648, Sigma; 1:200), acetylated tubulin (T6793; Sigma; 1:125), keratin (1h5; Developmental Studies Hybridoma Bank; 1:500), and GFP (N86/8; Developmental Studies Hybridoma Bank; 1:50) and incubated overnight on a nutator at 4 °C. After washing, the samples were incubated with the appropriate secondary antibody at a 1:200 dilution overnight at 4 °C. F-actin and cell nuclei were visualized with BODIPYFL–phallacidin (2.5:1000) and either YO-PRO (1:10000, Invitrogen) or Hoechst 33342 (1:2000, Thermo Fisher), respectively. Embryos were then bisected using forceps and a blade into transverse, sagittal, or en fas sections. These bisections were then dehydrated in an ethanol series and cleared in 2:1 benzyl benzoate:benzyl alcohol (BB:BA).

Fixed samples were imaged with an inverted compound microscope (Leica) with a 63x/1.40 NA oil immersion, a 25x/0.95 NA water immersion objective lens, or a 10x/0.95NA air objective lens equipped with a spinning disk scanhead (Yokogawa) and a CMOS camera (Hamamatsu). Sequential images were acquired using a microscope automation software (μManager 2.0)(Edelstein et al., 2014). For *en face* images, tiled images were stitched together using an ImageJ plugin (Horl et al., 2019). Planar images from 3D surfaces were extracted using SheetMeshProj (Wada and Hayashi, 2020).

### Live Imaging and Image Analysis

Brightfield time-lapses were collected using a phase-contrast microscope, Zeiss Axiovert S 100 (White Plains, NY, USA), equipped with a camera DMK23G445 (The ImagingSource, Charlotte, NC, USA), XY precision stage (Ludl, Hawthorne, NY), a Z-focus (Heidenhain, Traunreut, Germany) and a 10x objective. Sequential images were acquired using a microscope automation software (μManager 1.4)(Edelstein et al., 2014). Time-lapses of fluorescently labeled embryos were collected using a confocal laser scan head (SP5 Leica Microsystems) mounted on an inverted compound microscope (DMI6000, Leica Microsystems) equipped with a 10X air objective using acquisition software (LASAF, Leica Microsystems). See Supplemental Methods for Image Analysis Pipeline (Supplemental Material S1).

### Statistical Analysis

All statistical analyses were performed using GraphPad PRISM (v10.4.1), except for circular statistics, which were conducted in RStudio (R4.4.3). Normality was assessed using the Shapiro-Wilk test, and when the assumption of normality was violated, a Mann-Whitney U test was used for group comparisons. To evaluate the relationship between time and distance to the axis of rotation, a linear regression analysis was performed on tracked nuclear trajectories. Quadratic and logarithmic fits were also evaluated but did not substantially improve model performance or alter the interpretation. A Chi-Square goodness-of-fit test was applied to compare the frequency distribution of fiber orientations between treatment groups. Multiple embryos were used for all experimental replicates. Each experiment was conducted in at least three independent replicate clutches (embryos isolated from three separate frogs).

## Supplemental Information

Supplemental Material S1

Figures S1–S6 and supplemental references

Video S1. Posterior neuropore morphogenesis, related to Figure 1

Video S2. Nuclear dynamics in the posterior neuropore, related to Figure 2

Video S3. Membrane-labeled cells in the posterior neuropore, related to Figure 2

Video S4. Membrane-labeled cells in the lateral ectoderm, related to Figure 2

Video S5. Membrane-labeled cells in the ventral tissues, related to Figure 2

Video S6. Ventralized and dorsalized embryos, related to Figure 4

Video S7. Fibronectin disrupted embryos, related to Figure 6

## References

Anjum, S., Vijayraghavan, D., Fernandez-Gonzalez, R., Sutherland, A. and Davidson, L. (2025) ‘Inferring active and passive mechanical drivers of epithelial convergent extension’, bioRxiv: 2025.01.28.635314.

Asai, R., Prakash, V. N., Sinha, S., Prakash, M. and Mikawa, T. (2024) ‘Coupling and uncoupling of midline morphogenesis and cell flow in amniote gastrulation’, Elife 12: RP89948.

Banavar, S. P., Carn, E. K., Rowghanian, P., Stooke-Vaughan, G., Kim, S. and Campas, O. (2021) ‘Mechanical control of tissue shape and morphogenetic flows during vertebrate body axis elongation’, Sci Rep 11(1): 8591.

Beck, C. W. (2015) ‘Development of the vertebrate tailbud’, Wiley Interdiscip Rev Dev Biol 4(1): 33–44.

Briggs, J. A., Weinreb, C., Wagner, D. E., Megason, S., Peshkin, L., Kirschner, M. W. and Klein, A. M. (2018) ‘The dynamics of gene expression in vertebrate embryogenesis at single-cell resolution’, Science 360(6392): eaar5780.

Catala, M. (2021) ‘Overview of Secondary Neurulation’, J Korean Neurosurg Soc 64(3): 346–358.

Cetera, M., Leybova, L., Joyce, B. and Devenport, D. (2018) ‘Counter-rotational cell flows drive morphological and cell fate asymmetries in mammalian hair follicles’, Nat Cell Biol 20(5): 541–552.

Chitnis, A., Henrique, D., Lewis, J., Ish-Horowicz, D. and Kintner, C. (1995) ‘Primary neurogenesis in Xenopus embryos regulated by a homologue of the Drosophila neurogenic gene Delta’, Nature 375(6534): 761–6.

Chu, C. W. and Davidson, L. A. (2022) ‘Chambers for Culturing and Immobilizing Xenopus Embryos and Organotypic Explants for Live Imaging’, Cold Spring Harb Protoc 2022(5): Pdb prot107649.

Davidson, L. A. (2011) ‘Embryo mechanics: balancing force production with elastic resistance during morphogenesis’, Curr Top Dev Biol 95: 215–41.

Davidson, L. A. (2024) ‘Gears of life: A primer on the simple machines that shape the embryo’, Curr Top Dev Biol 160: 87–109.

Davidson, L. A., Keller, R. and DeSimone, D. W. (2004) ‘Assembly and remodeling of the fibrillar fibronectin extracellular matrix during gastrulation and neurulation in Xenopus laevis’, Dev Dyn 231(4): 888–95.

Davidson, L. A. and Keller, R. E. (1999) ‘Neural tube closure in Xenopus laevis involves medial migration, directed protrusive activity, cell intercalation and convergent extension’, Development 126(20): 4547–56.

Davidson, L. A., Marsden, M., Keller, R. and Desimone, D. W. (2006) ‘Integrin alpha5beta1 and fibronectin regulate polarized cell protrusions required for Xenopus convergence and extension’, Curr Biol 16(9): 833–44.

Edelstein, A. D., Tsuchida, M. A., Amodaj, N., Pinkard, H., Vale, R. D. and Stuurman, N. (2014) ‘Advanced methods of microscope control using muManager software’, J Biol Methods 1(2).

Firmino, J., Rocancourt, D., Saadaoui, M., Moreau, C. and Gros, J. (2016) ‘Cell Division Drives Epithelial Cell Rearrangements during Gastrulation in Chick’, Dev Cell 36(3): 249–61.

Gjorevski, N. and Nelson, C. M. (2010) ‘Endogenous patterns of mechanical stress are required for branching morphogenesis’, Integr Biol (Camb) 2(9): 424–34.

Gont, L. K., Steinbeisser, H., Blumberg, B. and de Robertis, E. M. (1993) ‘Tail formation as a continuation of gastrulation: the multiple cell populations of the Xenopus tailbud derive from the late blastopore lip’, Development 119(4): 991–1004.

Guillon, E., Das, D., Julich, D., Hassan, A. R., Geller, H. and Holley, S. (2020) ‘Fibronectin is a smart adhesive that both influences and responds to the mechanics of early spinal column development’, Elife 9: e48964.

Harris, W. A. and Hartenstein, V. (1991) ‘Neuronal determination without cell division in Xenopus embryos’, Neuron 6(4): 499–515.

Henkels, J., Oh, J., Xu, W., Owen, D., Sulchek, T. and Zamir, E. (2013) ‘Spatiotemporal mechanical variation reveals critical role for rho kinase during primitive streak morphogenesis’, Ann Biomed Eng 41(2): 421–32.

His, W. (1888) ‘On the principles of animal morphology.’, Proceedings of Royal Society of Edinburgh 15: 287–298.

Horl, D., Rojas Rusak, F., Preusser, F., Tillberg, P., Randel, N., Chhetri, R. K., Cardona, A., Keller, P. J., Harz, H., Leonhardt, H. et al. (2019) ‘BigStitcher: reconstructing high-resolution image datasets of cleared and expanded samples’, Nat Methods 16(9): 870–874.

Jackson, T. R., Kim, H. Y., Balakrishnan, U. L., Stuckenholz, C. and Davidson, L. A. (2017) ‘Spatiotemporally Controlled Mechanical Cues Drive Progenitor Mesenchymal-to-Epithelial Transition Enabling Proper Heart Formation and Function’, Curr Biol 27(9): 1326–1335.

Kao, K. R. and Elinson, R. P. (1988) ’The entire mesodermal mantle behaves as Spemann’s organizer in dorsoanterior enhanced Xenopus laevis embryos’, Dev Biol 127(1): 64–77.

Kao, K. R. and Elinson, R. P. (1989) ‘Dorsalization of mesoderm induction by lithium’, Dev Biol 132(1): 81–90.

Kato, S. and Inomata, H. (2023) ‘Blastopore gating mechanism to regulate extracellular fluid excretion’, iScience 26(5): 106585.

Keller, R. (2002) ‘Shaping the vertebrate body plan by polarized embryonic cell movements’, Science 298(5600): 1950–4.

Keller, Ray and Sutherland, Ann (2020) Convergent extension in the amphibian, Xenopus laevis Gastrulation: From Embryonic Pattern to Form, vol. 136.

Kinoshita, N., Sasai, N., Misaki, K. and Yonemura, S. (2008) ‘Apical accumulation of Rho in the neural plate is important for neural plate cell shape change and neural tube formation’, Mol Biol Cell 19(5): 2289–99.

Loganathan, R., Rongish, B. J., Smith, C. M., Filla, M. B., Czirok, A., Benazeraf, B. and Little, C. D. (2016) ‘Extracellular matrix motion and early morphogenesis’, Development 143(12): 2056–65.

Lowery, L. A. and Sive, H. (2004) ‘Strategies of vertebrate neurulation and a re-evaluation of teleost neural tube formation’, Mech Dev 121(10): 1189–97.

Masak, G. and Davidson, L. A. (2023) ‘Constructing the pharyngula: Connecting the primary axial tissues of the head with the posterior axial tissues of the tail’, Cells Dev 176: 203866.

Miao, Y. and Pourquie, O. (2024) ‘Cellular and molecular control of vertebrate somitogenesis’, Nat Rev Mol Cell Biol 25(7): 517–533.

Mitsui, Y. and Schneider, E. L. (1976) ‘Relationship between cell replication and volume in senescent human diploid fibroblasts’, Mech Ageing Dev 5(1): 45–56.

Mongera, A., Rowghanian, P., Gustafson, H. J., Shelton, E., Kealhofer, D. A., Carn, E. K., Serwane, F., Lucio, A. A., Giammona, J. and Campas, O. (2018) ‘A fluid-to-solid jamming transition underlies vertebrate body axis elongation’, Nature 561(7723): 401–405.

Morita, H., Kajiura-Kobayashi, H., Takagi, C., Yamamoto, T. S., Nonaka, S. and Ueno, N. (2012) ‘Cell movements of the deep layer of non-neural ectoderm underlie complete neural tube closure in Xenopus’, Development 139(8): 1417–26.

Myers, C. T., Appleby, S. C. and Krieg, P. A. (2014) ‘Use of small molecule inhibitors of the Wnt and Notch signaling pathways during Xenopus development’, Methods 66(3): 380–9.

Newman, K., Aguero, T. and King, M. L. (2018) ‘Isolation of Xenopus Oocytes’, Cold Spring Harb Protoc 2018(2).

Nieuwkoop PD, Faber J. (1956) Normal table of Xenopus laevis (Daudin). A systematical and chronological survey of the development from the fertilized egg till the end of metamorphosis, Amsterdam, The Netherlands: North-Holland Publishing Company.

Palmquist, K. H., Tiemann, S. F., Ezzeddine, F. L., Yang, S., Pfeifer, C. R., Erzberger, A., Rodrigues, A. R. and Shyer, A. E. (2022) ‘Reciprocal cell-ECM dynamics generate supracellular fluidity underlying spontaneous follicle patterning’, Cell 185(11): 1960–1973 e11.

Pokrovsky, D., Forne, I., Straub, T., Imhof, A. and Rupp, R. A. W. (2021) ‘A systemic cell cycle block impacts stage-specific histone modification profiles during Xenopus embryogenesis’, PLoS Biol 19(9): e3001377.

Rauzi, M. (2020) ‘Cell intercalation in a simple epithelium’, Philos Trans R Soc Lond B Biol Sci 375(1809): 20190552.

Roca-Cusachs, P., Gauthier, N. C., Del Rio, A. and Sheetz, M. P. (2009) ‘Clustering of alpha(5)beta(1) integrins determines adhesion strength whereas alpha(v)beta(3) and talin enable mechanotransduction’, Proc Natl Acad Sci U S A 106(38): 16245–50.

Rozario, T. and DeSimone, D. W. (2010) ‘The extracellular matrix in development and morphogenesis: a dynamic view’, Dev Biol 341(1): 126–40.

Saadaoui, M., Rocancourt, D., Roussel, J., Corson, F. and Gros, J. (2020) ‘A tensile ring drives tissue flows to shape the gastrulating amniote embryo’, Science 367(6476): 453–458.

Sato, K., Hiraiwa, T., Maekawa, E., Isomura, A., Shibata, T. and Kuranaga, E. (2015) ‘Left-right asymmetric cell intercalation drives directional collective cell movement in epithelial morphogenesis’, Nat Commun 6(1): 10074.

Scott, L. E., Mair, D. B., Narang, J. D., Feleke, K. and Lemmon, C. A. (2015) ‘Fibronectin fibrillogenesis facilitates mechano-dependent cell spreading, force generation, and nuclear size in human embryonic fibroblasts’, Integr Biol (Camb) 7(11): 1454–65.

Shawky, J. H. and Davidson, L. A. (2015) ‘Tissue mechanics and adhesion during embryo development’, Dev Biol 401(1): 152–64.

Shook, D. R., Wen, J. W. H., Rolo, A., O’Hanlon, M., Francica, B., Dobbins, D., Skoglund, P., DeSimone, D. W., Winklbauer, R. and Keller, R. E. (2022) ’Characterization of convergent thickening, a major convergence force producing morphogenic movement in amphibians’, Elife 11: e57642.

Singh, P., Carraher, C. and Schwarzbauer, J. E. (2010) ‘Assembly of fibronectin extracellular matrix’, Annu Rev Cell Dev Biol 26(Volume 26, 2010): 397–419.

Sinha, P., Sarkar, K., Pandey, B. and Nandi, Nityananda (2013) ‘Experimental Study on Vortex Motion’, International Journal of Fluid Mechanics Research 40(6): 512–519.

Skoglund, P. M. (1996) What Xenopus fibrillin does in the early embryo; clues from a dominant negative approach. United States -- California: University of California, San Diego.

Sokol, S. Y. (1996) ‘Analysis of Dishevelled signalling pathways during Xenopus development’, Curr Biol 6(11): 1456–67.

Stooke-Vaughan, G. A., Kim, S., Yen, S. T., Son, K., Banavar, S. P., Giammona, J., Kimelman, D. and Campas, O. (2025) ‘The physical roles of different posterior tissues in zebrafish axis elongation’, Nat Commun 16(1): 1839.

Stubbs, J. L., Davidson, L., Keller, R. and Kintner, C. (2006) ‘Radial intercalation of ciliated cells during Xenopus skin development’, Development 133(13): 2507–15.

Tao, Q., Yokota, C., Puck, H., Kofron, M., Birsoy, B., Yan, D., Asashima, M., Wylie, C. C., Lin, X. and Heasman, J. (2005) ‘Maternal wnt11 activates the canonical wnt signaling pathway required for axis formation in Xenopus embryos’, Cell 120(6): 857–71.

Tian, S. L., Gao, Y. S., Dong, X. R. and Liu, C. Q. (2018) ‘Definitions of vortex vector and vortex’, Journal of Fluid Mechanics 849: 312–339.

Ventura, G. and Sedzinski, J. (2022) ‘Emerging concepts on the mechanical interplay between migrating cells and microenvironment in vivo’, Front Cell Dev Biol 10: 961460.

Wada, H. and Hayashi, S. (2020) ‘Net, skin and flatten, ImageJ plugin tool for extracting surface profiles from curved 3D objects’, MicroPubl Biol 2020.

Wallingford, J. B. and Harland, R. M. (2002) ‘Neural tube closure requires Dishevelled-dependent convergent extension of the midline’, Development 129(24): 5815–25.

Wang, J. X. and White, M. D. (2021) ‘Mechanical forces in avian embryo development’, Semin Cell Dev Biol 120: 133–146.

Wang, X., Merkel, M., Sutter, L. B., Erdemci-Tandogan, G., Manning, M. L. and Kasza, K. E. (2020) ‘Anisotropy links cell shapes to tissue flow during convergent extension’, Proc Natl Acad Sci U S A 117(24): 13541–13551.

Wu, D., Yamada, K. M. and Wang, S. (2023) ‘Tissue Morphogenesis Through Dynamic Cell and Matrix Interactions’, Annu Rev Cell Dev Biol 39(1): 123–144.

Xiong, F., Ma, W., Benazeraf, B., Mahadevan, L. and Pourquie, O. (2020) ‘Mechanical Coupling Coordinates the Co-elongation of Axial and Paraxial Tissues in Avian Embryos’, Dev Cell 55(3): 354–366 e5.

Zahn, N., James-Zorn, C., Ponferrada, V. G., Adams, D. S., Grzymkowski, J., Buchholz, D. R., Nascone-Yoder, N. M., Horb, M., Moody, S. A., Vize, P. D. et al. (2022) ‘Normal Table of Xenopus development: a new graphical resource’, Development 149(14).

Zamir, E. A., Rongish, B. J. and Little, C. D. (2008) ‘The ECM moves during primitive streak formation--computation of ECM versus cellular motion’, PLoS Biol 6(10): e247.

Zitong, C., Zhenyu, C. and Rinkevich, Y. (2024) ‘Tissue fluidity: biophysical shape-shifting for regeneration’, Signal Transduct Target Ther 9(1): 329.

