## Supplemental Figures and Methods for "Supracellular Mechanics and Counter-Rotational Bilateral Flows Orchestrate Posterior Morphogenesis"

**Table of Contents - Supplementary Materials:**

**I. Supplementary Methods.**

**II. Supplementary Figures.**

**III. Supplementary Videos.**

### I. Supplementary Methods.

**Rotational Analysis:** To quantify the degree of rotational motion using vorticity, a fundamental measure in fluid dynamics that describes the local rotation of a velocity field (Asai et al., 2024). Vorticity ($\omega$) is defined mathematically as the curl of the velocity field:

$$\omega = \nabla \times v \qquad \text{Equation 1}$$

where $v$ represents the velocity vector field. To quantify the degree of rotational motion within posterior tissues, we calculated vorticity from velocity maps:

$$\omega = \frac{\partial v_y}{\partial x} - \frac{\partial v_x}{\partial y} \qquad \text{Equation 2}$$

where $v_x$ and $v_y$ represent the velocity components in the x and y directions, respectively. Higher vorticity values correspond to stronger rotational forces, while the sign of vorticity distinguishes clockwise ($\omega<0$) and counterclockwise ($\omega>0$) rotation. Because vorticity is a spatial derivative of velocity, it provides a direct measure of localized rotational motion, independent of translational movements.

**Axial Strain Analysis:** To quantify deformation along the anteroposterior axis we calculated axial strain ($\epsilon_{yy}$). This strain represents material deformation by measuring the local extension or compression along the anterior-posterior axis (here the y-axis). We used the following equation to calculate axial strain from velocity maps:

$$\varepsilon_{yy} = \frac{\partial v_y}{\partial y} \qquad \text{Equation 3}$$

where $v_y$ is the velocity component along the y-direction. This equation represents the rate of change of velocity in the y-direction, indicating how much the material is stretching ($\epsilon_{yy} > 0$) or compressing ($\epsilon_{yy} < 0$).

**SWIRL Analysis:** To further characterize rotational movements we adapted the SWIRL vortex detection algorithm developed to quantify vortices and rotational movement in the solar atmosphere (Canivete Cuissa and Steiner, 2024). We were able to filter out vortices that exhibited characteristics of shear-dominated flow and provide quantitative metrics to describe vortex properties (Fig 1C,G). The first of these measures, swirling strength (S), quantifies the intensity of rotational motion by capturing the imaginary component of the velocity gradient tensor's eigenvalues. Swirling strength is given by:

$$S = 2 \mid Im(\lambda) \mid \qquad \text{Equation 4}$$

where λ represents the complex eigenvalues of the velocity gradient tensor, and *Im(λ)* denotes its imaginary part, which corresponds to local rotational flow. Swirling strength is high when vortices are well-defined and persistent over time. The second measure, compactness (ζ), assesses the spatial coherence of vortex structures by evaluating the ratio of shear to rotational motion. Compactness is given by:

$$\zeta = \frac{Re(\lambda)}{Im(\lambda)} \qquad \text{Equation 5}$$

Where *Re(λ)* represents the real component of the eigenvalue, which reflects shear-driven deformation. Low compactness values indicate that rotational motion dominates over shear deformation, meaning the vortex is spatially constrained rather than diffuse or transient.

To distinguish changes in the overall symmetry of vortices, we also compared the swirling strength of vortices found in the same embryo, calculating the Vortex Asymmetry Index (VAI) as:

$$VAI = \frac{2|S_1 - S_2|}{S_1 + S_2} \quad \text{Equation 6}$$

where $S_1$ and $S_2$ represent the swirling strength values of the two vortices within the same embryo, $|S_1 - S_2|$ computes the absolute difference in swirling strength, and the factor of 2 normalizes the asymmetry relative to the total swirling strength of both vortices. Lower values of VAI would indicate greater symmetry between vortices with VAI = 0 reflecting perfectly symmetrical vortices, while greater VAI (VAI > 0) indicates asymmetric vortices.

**Nuclear Segmentation and Tracking:** Maximum-intensity projections of H2B-mScarlet confocal timelapse Z-stacks were generated prior to segmentation and tracking. Image stacks were pre-processed in FIJI to improve segmentation and tracking accuracy. Nuclear segmentation was performed using either StarDist 2D (Schmidt et al., 2018) or Cellpose (Stringer et al., 2021), depending on image quality, followed by tracking with TrackMate (Ershov et al., 2022). Segmentation and tracking parameters were optimized for each dataset, and spots and tracks were filtered using custom thresholds. Nuclei were linked across frames using the Advanced Kalman Tracker. After automated filtering, all tracks were reviewed manually and erroneous detections were corrected or removed to ensure consistency across samples.

The center of rotation was estimated from maximum-intensity projections across all time points, with regions of interest (ROIs) manually selected at the spiral centers. For each nucleus, the distance from the nearest rotational center was calculated in TrackMate using the ROI set (Fig.

2A–B). Tracks with an average distance greater than 300 μm from the rotational center were excluded from analysis to restrict measurements to nuclei within the vortex.

**Membrane Segmentation and Tracking:** Maximum-intensity projections of CMV:memGFP confocal timelapse Z-stacks were generated prior to segmentation and tracking. To improve segmentation accuracy, images were pre-processed in FIJI with background subtraction to reduce noise and enhance membrane contrast. Whole-body rotational drift was corrected by rigid-body registration using the Linear Stack Alignment with SIFT plugin (Lowe, 2004).

Segmentation was performed using Cellpose (Stringer et al., 2021), and tracks were generated with TrackMate (Ershov et al., 2022) following the nuclear segmentation workflow. Tracks corresponding to dividing or intercalating cells were excluded by filtering out trajectories without a corresponding cell in every frame. To account for variability in aspect ratios across cells and differences in imaging durations among replicates, aspect ratios were normalized in Excel, using each cell's initial aspect ratio as the reference.

**T1 Transition Analysis:** We tracked T1 transitions, in which four neighboring cells rearrange through junction shrinkage and neighbor exchange (Fig S2A-C). To assess how much time cells spend in a T1 transition state over two hours, we calculated the T1 dwell fraction, defined as the proportion of total contact time spent in a T1 state:

$$\text{T1 dwell fraction} = \frac{\sum T1_{time}}{Total\ time} \qquad \text{Equation 7}$$

where $T1_{time}$ represents the time a cell spends in a T1 transition, from entrance to exit. Higher T1 dwell fraction values indicate that cells remain in T1 transitions for a larger proportion of time, reflecting more persistent junctional rearrangements.

To quantify the duration of each completed T1 transition, we computed the mean T1 duration, defined as:

$$\text{Mean T1 duration} = \frac{1}{N_j}\sum_{i=1}^{N_j} T1_{duration,i,j} \qquad \text{Equation 8}$$

where $j$ is the index of each completed T1 transition, $N_j$ is the number of cells that participated in the $j$-th T1 transition, and $T1_{duration,i,j}$ represents the duration of the $j$-th T1 transition for cell $i$, measured as the number of consecutive time points the cell remained in a T1 state before completing the transition.

**Fiber Analysis:** To quantify fibronectin organization, maximum-intensity projections were generated from linearized image stacks (Wada and Hayashi, 2020), selectively projecting only the superficial layer to separate superficial from deep fibronectin networks and minimize staining artifacts (Fig. S4A-B). For consistent quantification, 100 × 100 μm regions of interest were cropped from areas at least 100 μm ventral to the blastopore. Images were pre-processed in FIJI using Contrast Limited Adaptive Histogram Equalization (CLAHE) (Zuiderveld, 1994), despeckling, and outlier removal to enhance fiber visibility and correct for uneven illumination.

Fiber morphology was quantified using DiameterJ (Hotaling et al., 2015), which calculates fiber diameter, pore size, and intersection density (Fig. S4C). To improve thresholding, DiameterJ was modified to incorporate an additional filtering step. In low-density images, inversion was applied to ensure accurate thresholding and skeletonization. Images with insufficient fiber density or disconnected networks were excluded, as pore analysis requires a minimum number of pores.

To quantify fiber orientation and assess global alignment, orientation distributions obtained from DiameterJ were binned into 15° intervals and plotted as polar histograms using Origin (v2025; OriginLab, Northampton, MA).

**Ciliated Cell Counting**: To quantify ciliated cell density, Z-stacks of tissues immunostained for acetylated tubulin were acquired and converted to maximum-intensity projections within a 250 µm × 250 µm region of interest. To improve segmentation accuracy, projected images were pre-processed with smoothing and Gaussian blur filters in FIJI. Automated segmentation and cell counting were performed using StarDist 2D (Schmidt et al., 2018), which generated a list of Regions of Interest (ROIs). ROIs were subsequently filled with color and reviewed manually to verify labeling accuracy (Fig S5B,D).

References

Asai, R., Prakash, V. N., Sinha, S., Prakash, M. and Mikawa, T. (2024) 'Coupling and uncoupling of midline morphogenesis and cell flow in amniote gastrulation', *Elife* 12: RP89948.

Canivete Cuissa, J. R. and Steiner, O. (2024) 'Innovative and automated method for vortex identification', *Astronomy & Astrophysics* 682.

Ershov, D., Phan, M. S., Pylvanainen, J. W., Rigaud, S. U., Le Blanc, L., Charles-Orszag, A., Conway, J. R. W., Laine, R. F., Roy, N. H., Bonazzi, D. et al. (2022) 'TrackMate 7: integrating state-of-the-art segmentation algorithms into tracking pipelines', *Nat Methods* 19(7): 829-832.

Hotaling, N. A., Bharti, K., Kriel, H. and Simon, C. G., Jr. (2015) 'DiameterJ: A validated open source nanofiber diameter measurement tool', *Biomaterials* 61: 327-38.

Lowe, D. G. (2004) 'Distinctive image features from scale-invariant keypoints', *International Journal of Computer Vision* 60(2): 91-110.

Mitsui, Y. and Schneider, E. L. (1976) 'Relationship between cell replication and volume in senescent human diploid fibroblasts', *Mech Ageing Dev* 5(1): 45-56.

Myers, C. T., Appleby, S. C. and Krieg, P. A. (2014) 'Use of small molecule inhibitors of the Wnt and Notch signaling pathways during Xenopus development', *Methods* 66(3): 380-9.

Schmidt, U., Weigert, M., Broaddus, C. and Myers, G. (2018) 'Cell Detection with Star-Convex Polygons', *Lecture Notes in Computer Science*.

Stringer, C., Wang, T., Michaelos, M. and Pachitariu, M. (2021) 'Cellpose: a generalist algorithm for cellular segmentation', *Nat Methods* 18(1): 100-106.

Wada, H. and Hayashi, S. (2020) 'Net, skin and flatten, ImageJ plugin tool for extracting surface profiles from curved 3D objects', *MicroPubl Biol* 2020.

Zuiderveld, K. (1994) 'Contrast limited adaptive histogram equalization', *Graphics gems IV*: 474-485.

### II. Supplementary Figures.

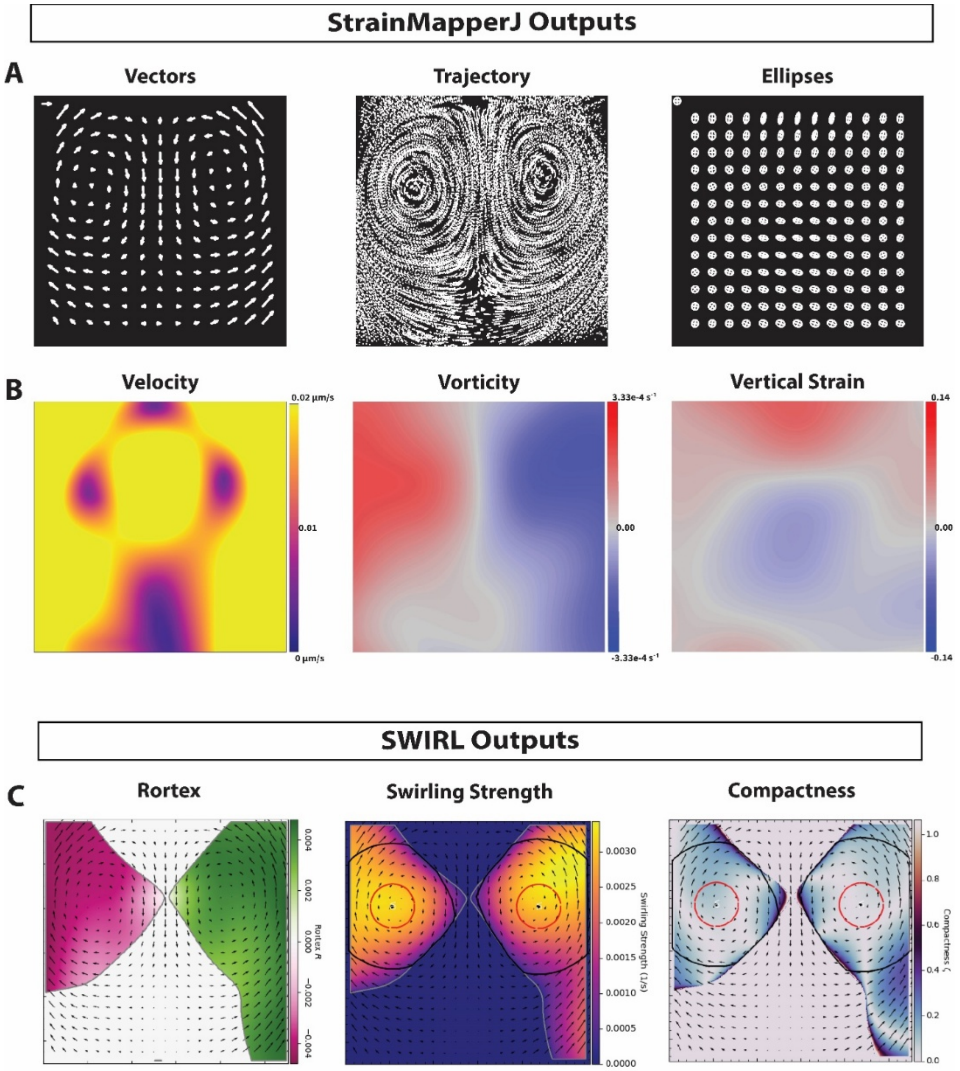


**Figure S1. Raw StrainMapperJ and SWIRL outputs used for kinematic analysis of posterior tissue movements.** (A-B) StrainMapperJ-generated maps from a representative dataset. (A) Vector and Ellipse maps from a single, representative timepoint. Each symbol is normalized to the maximum value within that timepoint. Trajectory maps display the cumulative displacement of randomly distributed dots, deformed over time to visualize tissue flow. (B) Velocity, vorticity, and vertical strain maps, averaged across all time-points, representing global kinematic trends across the imaging period. (C) SWIRL-generated vortex analysis maps. Rortex values, superimposed onto a velocity vector map, are used to identify coherent rotational flow structures. Red circles indicate the area from which swirling strength and compactness are measured, providing a basis for comparisons between vortices and across experimental treatments. Black circles represent the estimated vortex boundary determined by the SWIRL algorithm. Related to Figure 1.

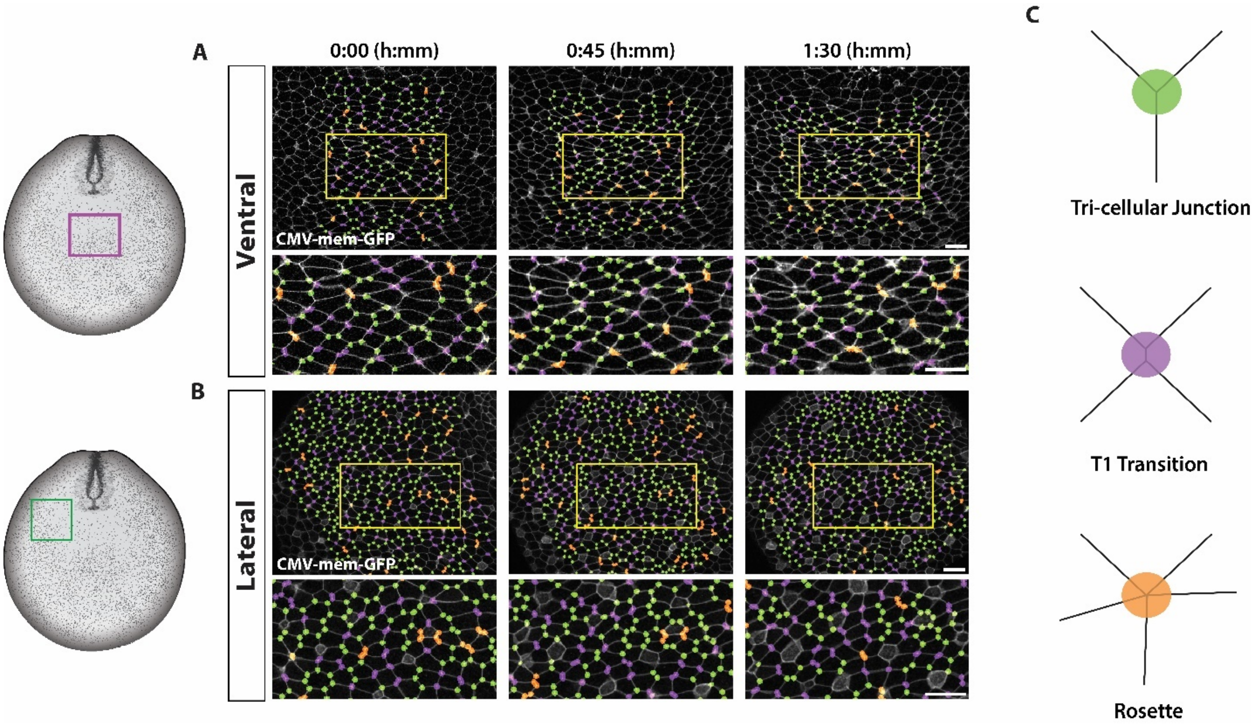


**Figure S2. Junction analysis reveals abundant T1 and Rosettes in membrane-labeled embryos** (A-B) Graphic illustration of a stage 18 embryo, indicating the approximate imaging region (box) relative to the blastopore. (A) Vertex analysis of the ventral region from registered timelapse series of CMV-mem-GFP embryos. A magnified view highlights the number of cells associated with junctions during lateral cell movements in ventral tissues. (B) Vertex analysis of the lateral region from registered timelapse series of CMV-mem-GFP embryos. A magnified view highlights the number of cells associated with junctions during tissue rotation in side tissues. (C) Diagram representing vertex classification and color schema. Tri-cellular junctions (3-cell vertices) are represented by green circles. T1 transitions (4-cell vertices) are represented by purple circles. Rosettes (>5-cell vertices) are represented by orange circles. Scale bars: 50 μm in A-B. Related to Figure 2.

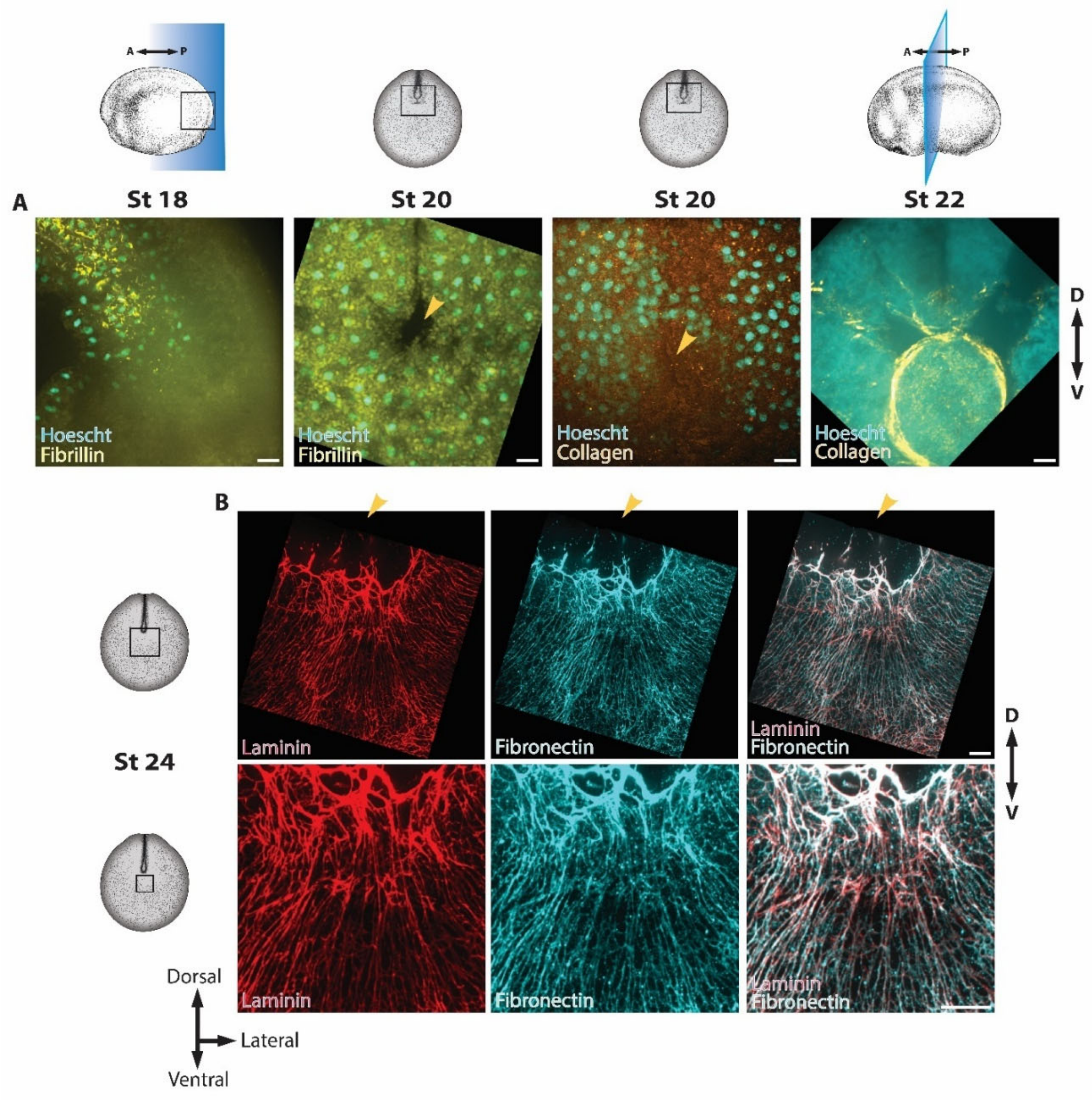


**Figure S3. Fibronectin and Laminin are the predominant ECM components in posterior tissues.** (A-B) Schematic illustrations of embryos at the indicated stages, with boxes marking the approximate imaging regions relative to the blastopore. (A) Immunostained images of embryos labeled with antibodies against Fibrillin, Collagen, and Hoechst from stages 18–22. Sagittal section of the posterior neural tube shows Fibrillin fibers localized in the presomitic mesoderm, tapering off posteriorly. En face views of the posterior-most tissues reveal an absence of both Fibrillin and Collagen surrounding the blastopore (orange arrowhead). A transverse section highlights Collagen localization around the notochord. Blue shaded boxes indicate the imaging planes for sagittal and transverse sections. (B) Immunostained images of stage 24 embryos labeled with antibodies against Laminin and Fibronectin. Magnified views of the region ventral to the blastopore (orange arrowhead) show co-localization of Laminin and Fibronectin. Scale bars: 20 µm in A-B. Related to Figure 3.

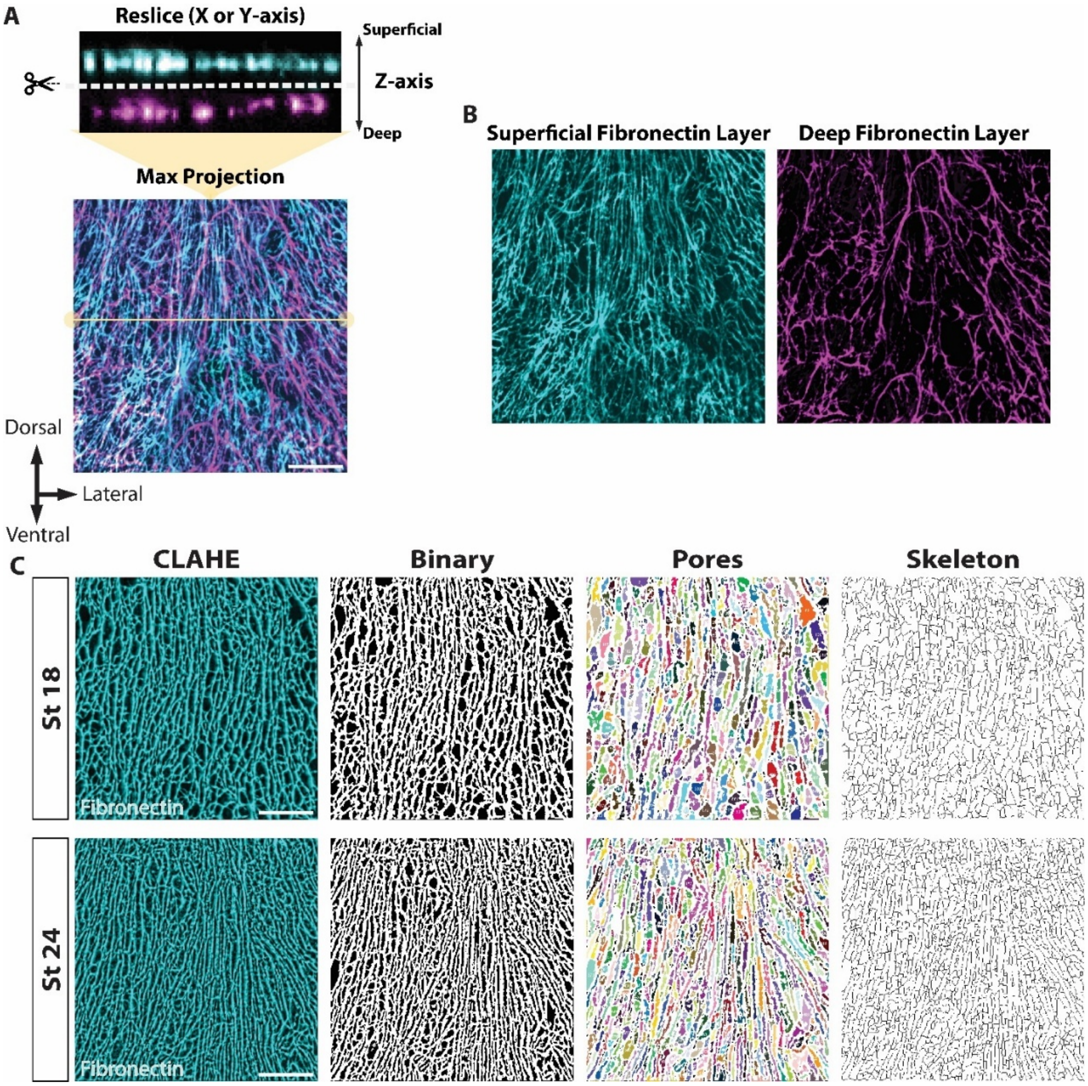


**Figure S4. Layer-Specific Analysis of Fibronectin Organization Using Z-Stack Linearization and DiameterJ Quantification.** (A) Reslice of immunostained Z-stack images stained with an antibody against fibronectin, processed with CLAHE to enhance fiber contrast and correct uneven illumination. Lookup tables are applied to distinguish the superficial and deep fibronectin layers. A yellow line on the max-projected image represents the location of the reslice. A dotted line with scissors indicates the separation of superficial and deep fibronectin layers for further analysis (B) CLAHE-processed max-projected images of superficial and deep fibronectin layers after linearization using SheetMeshProj (Wada and Hayashi, 2020), enabling layer-specific visualization of fiber organization. (C) CLAHE-processed immunostained fibronectin images serve as input for binarization, which is used to generate skeletons, measure fiber density, and identify pores. Binarized images are overlaid with identified pore regions, and extracted skeletons are used to calculate fiber intersection density and diameter. Images in (C) were further processed with CLAHE and noise removal to improve binarization accuracy. Scale bars: 20 µm in A,C. Related to Figure 3.

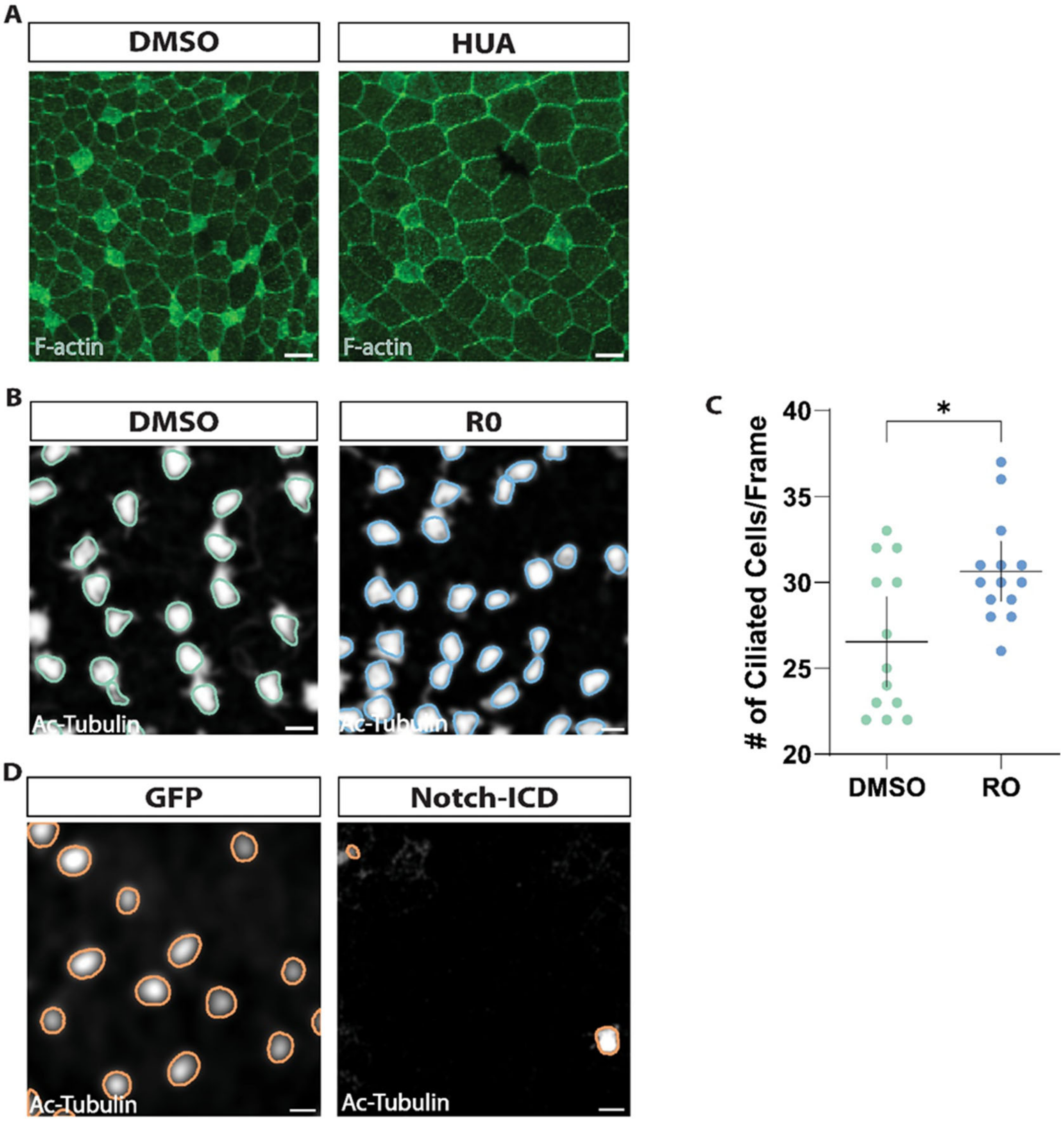


**Figure S5. Alterations in cell size and ciliated cell density following HUA, R0, and Notch-ICD treatments.** (A-B) Max-projected images of immunostained ventral tissues. (A) En face sections of stage 26 embryos treated with DMSO and HUA, stained against F-actin to assess changes in cell size as a validation of cell division inhibition (Mitsui and Schneider, 1976). (B-C) En face sections of posterior tissues immunostained against α-Tubulin to assess cilia density (Myers et al., 2014). Samples were processed with de-noising and Gaussian blur methods to enhance segmentation of cilia. Colored ROIs highlight cilia identified by StarDist. (B) R0-treated embryos exhibit an increased density of ciliated cells. (C) Quantification of ciliated cells in samples treated with DMSO and R0 within a 250μm x 250μm region. Each symbol represents the total number of cilia counted per embryo. Statistical significance was assessed using a Mann-Whitney U test (∗p=0.0333). Bars; mean ± 95% CI. (D) Notch-ICD-injected embryos show a reduced number of ciliated cells compared to GFP-injected controls (3 embryos per treatment). Scale bars: 20 μm in A-C. Related to Figure 5.

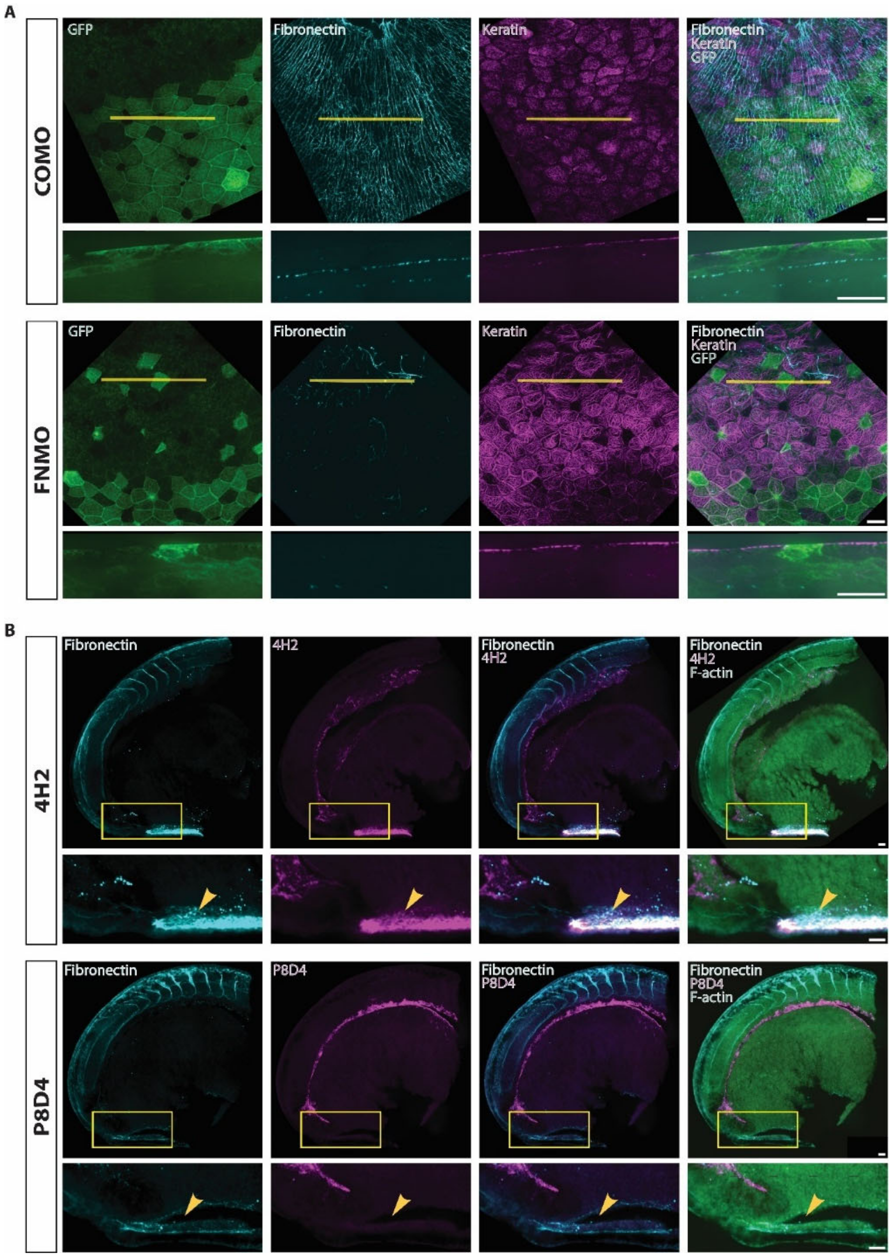


**Figure S6. Disruptions in ECM organization and tissue integrity following P8D4 and FNMO treatments.** (A-B) Max-projected images of immunostained posterior tissues. (A) En fas sections of posterior tissues in stage 20 embryos injected with morpholinos and 3X-GFP mRNA, and immunostained against GFP, Fibronectin, and Keratin. Z-stack images were linearized to isolate fibronectin layers before maximum intensity projection. XZ re-slices (yellow line) were generated from Z-stack images to visualize fibronectin distribution across tissue layers, revealing a reduction of fibronectin in FNMO-injected embryos. (B) Sagittal sections of posterior tissues in stage 22 embryos injected with 4H2 or P8D4 antibodies and immunostained against F-action, Fibronectin, and Keratin. Magnified view (yellow box) of ventral endoderm which is detached from superficial cell layers in P8D4 injected samples (orange arrow). Scale bars: 20 μm in A-B. Related to Figure 6.

### III. Supplementary Videos.

**Supplemental Videos S1–S7.** Live imaging and quantitative kinematic analyses of posterior morphogenesis in Xenopus laevis. Videos show representative brightfield and confocal timelapses, with overlays illustrating nuclear trajectories, cell behaviors, and tissue deformation maps. Together, these data highlight counter-rotational tissue flows, cell- and tissue-scale remodeling, and the effects of experimental perturbations on posterior morphogenesis.

**Video S1. Brightfield timelapse of posterior neuropore morphogenesis.** Embryo imaged from stage 18 to 20, corresponding to Fig. 1D, with cumulative tissue displacement mapped by a yellow hexagonal mesh overlay. Left, processed brightfield timelapse without overlay; right, same timelapse with mesh overlay indicating cell/pixel displacements.

**Video S2. Nuclear tracking reveals dynamic trajectories in the posterior neuropore.** Spinning disk confocal timelapse of an H2B-mScarlet–labeled embryo from stage 18. TrackMate-based segmentation and tracking shows nuclei color-coded by distance from the center of rotation (dot colors) and average nuclear velocity (track colors). Related to Fig. 2A.

**Video S3. Membrane dynamics at the blastopore lip during neural fold convergence.** Spinning disk timelapse of the posterior neuropore in a CMV-memGFP embryo. Cells located below the blastopore opening constrict while neural folds zipper together. Related to Fig. 2C.

**Video S4. Rotational tissue dynamics in lateral ectoderm.** Spinning disk timelapse of the posterior neuropore highlighting clockwise rotation of lateral tissues. A subset of tracked cells is outlined and color-coded by mean directional rate change (radians/sec). The blastopore exits the field of view as rotation progresses, and T1 transitions are observed throughout the tissue. Related to Fig. 2D.

**Video S5. Cell shape remodeling within ventral tissues.** Spinning disk timelapse of membrane-labeled embryos showing ventral epithelial cells outlined and color-coded by aspect ratio. The blastopore enters the field of view in the final frames. Related to Fig. 2E.

**Video S6. Flow dynamics in ventralized and dorsalized embryos.** Brightfield timelapse sequences of posterior tissues from stage 18. Top, processed brightfield images without overlays; bottom, corresponding deformation maps with yellow hexagonal mesh overlays. Panels are arranged left to right as labeled (dorsalized, control, ventralized). Related to Fig. 4C.

**Video S7. Posterior flow dynamics following antibody and morpholino perturbations.** Brightfield timelapse sequences of posterior tissues from stage 18. Top, processed brightfield images without overlays; bottom, corresponding deformation maps with yellow hexagonal mesh overlays. Panels are arranged left to right as labeled (4H2, P8D4, COMO, FNMO). Related to Fig. 6E.
